# Do we perform as well as we think we do? A systematic scoping review of self-evaluation of upper-extremity motor performance

**DOI:** 10.1101/2022.10.31.514569

**Authors:** Lucas D. Crosby, Gabriela Rozanski, Mira Browne, Avril Mansfield, Kara K. Patterson

## Abstract

The ability to self-evaluate motor performance or estimate performance errors is beneficial for motor learning or relearning in the context of neurologic injury. Some evidence suggests those with injury like stroke may be unable to accurately self-evaluate their performance; however, it is unclear if individuals who are absent of injury are accurate in this domain. We aimed to investigate the accuracy of self-evaluation and potential influencing factors by conducting a systematic search to identify literature involving the self- and objective-evaluation of upper-extremity motor tasks. Twenty-three studies satisfied inclusion criteria. Data revealed a moderate positive correlation between self- and objective evaluations across a variety of tasks, from trivial button pressing to specialized surgical suturing. Both under- and overestimation of performance was found across the papers. Key factors identified to influence the accuracy of self-evaluation were the task purpose, familiarity, difficulty, and whether an individual received a demonstration. This review identified some limitations in this field of research. Most notably, we found that very few studies have investigated the accuracy of self-evaluation of motor performance with the primary goal of comparison to objective performance. Many studies reported the data but did not make direct statistical comparisons. Moreover, due to inconsistencies between how self and objective-evaluations were conducted, we argue that in this area of investigation self-evaluation tools need to replicate the objective evaluation method, or at minimum the self-evaluation tool should ask questions specific to the construct of performance that is being measured objectively.

## Introduction

Precise and accurate completion of motor tasks is necessary for success during performance of activities in our everyday lives from simple tasks such as walking or reaching for a coffee cup to highly specialized and practiced technique like a baseball player swinging a bat, or a surgical resident suturing a wound to repair damage after injury. Investigating factors that influence successful task performance is essential to understand how people learn new motor skills, which may provide insights for the motor relearning process in rehabilitation. Some evidence demonstrates that if an individual is told to subjectively evaluate their movement performance throughout the acquisition phase, they perform better in retention tests compared to those who are not instructed to subjectively evaluate their movements (Guadagnoli & Kohl, 2001; Liu & Wrisberg, 1997). Self-evaluation of motor performance is thought to increase the clarity of the performance outcome (understanding why a certain result was achieved) and improve error-estimation capability (the ability to accurately detect intrinsic errors in performance) (Liu & Wrisberg, 1997).

Individuals must have accurate perception of their movement, to be able to effectively self-evaluate and apply feedback to improve motor performance. However, it has been shown that some individuals may not have optimal awareness regarding movement. In an experiment that obscured an arm movement error from vision, while the extrinsic feedback provided to the participant indicated that the desired outcome of the movement was achieved, the participants failed to report the movement error (Fourneret & Jeannerod, 1998). Therefore, an individual’s interpretation about the success of their movement from external sources may be prioritized over intrinsic sensory feedback about errors within the movement (Blakemore et al., 2002). Accurate self-evaluation of movement is difficult if an individual thinks they are performing better (or worse) than they truly are.

Understanding the accuracy of self-evaluation of motor performance has important application to the rehabilitation of neurological conditions such as stroke. Recovery or remediation of motor skills that may have been altered from damage often involves task repetition and correcting errors through therapists’ or other external feedback. However, it is not clear whether individuals are able to recognize errors in their own performance without extrinsic feedback. It is possible self-evaluation of motor performance could be disrupted by neurologic injury that impacts somatosensory pathways or higher-level processes like attention or awareness. However, an important first step is to understand this ability in intact or uninjured brains. Thus, the primary objective of this study was to investigate the nature and accuracy of self-evaluation of upper-extremity motor performance in healthy individuals. The secondary objective of this review was to determine factors that may influence motor performance self-evaluation.

## Methods

To achieve our objective, a systematic search of studies that had measured both objective motor performance and self-evaluation of a motor task performed by healthy adults was conducted. This scoping review was conducted following the Preferred Reporting Items for Systematic Reviews and Meta-Analyses (PRIMSA) guidelines for the scoping review process (Moher et al., 2009). Data extraction and charting was performed under the guidance of the Joanna Briggs Institute for the conduct of systematic scoping reviews (Peters et al., 2015).

### Literature search

An electronic search of the literature was conducted by an information specialist. Keywords and headings related to upper-extremity, motor tasks, and evaluation were identified prior to initiating the search. The following online databases were searched: CENTRAL, EMBASE, MEDLINE and MEDLINE in Process, PsychINFO, Pubmed, and SCOPUS. The initial search was conducted on January 11, 2017. Supplemental searches using the same strategy were conducted on August 23, 2018, and January 29, 2021. Three authors (LC, GR, MB) hand searched the reference list of the articles identified for inclusion and review articles that were identified as relevant during full-text review. A summary of the complete search strategy can be found in Appendix 1.

### Study selection

For inclusion, studies included: (i) a motor task of the upper extremity; (ii) participants’ self-evaluation of their performance on the task after performing it; (iii) an objective measurement of participants’ task performance as comparison; and (iv) healthy adult participants. Non-English studies, conference proceedings, and theses were excluded. In pairs, three reviewers (LC, GR, MB) screened titles and abstracts, and subsequently full texts, following the same procedures. Disagreements throughout the selection processes were resolved through discussion and consensus.

### Data extraction and analysis

In accordance with scoping review extracting and results charting guidelines (Peters et al., 2015), studies were qualitatively described. One author (LC or MB) extracted data from all included articles using a study-specific spreadsheet. Extracted data was checked for accuracy by a second author (GR or LC). Data extracted included study design, sample and subject characteristics, performance evaluation and analytical methods used, and key findings with respect to the relationship between self-evaluation and objective evaluation of task performance. Discrepancies in data extraction were resolved through discussion and consensus. Corresponding authors were contacted for additional information if the relationship between self-evaluation and objective evaluation was unclear.

Analytical method of comparison between self- and objective-evaluations of performance was extracted from the included articles along with statistical values (e.g., Pearson’s r, Cohen’s k) and significance level (p value). Additionally, the magnitude and direction of the discrepancy between performance and self-evaluation measures (i.e., degree of over- or under-estimation) was noted where possible in a descriptive summary of the data. This analysis was possible if two criteria were met: 1) the study reported a correlation between self- and objective-evaluation measures, and 2) the study reported individual scores for each measure or the mean scores for self- and objective-evaluation of performance. Over-estimation of performance indicates that the subjects believed they performed better (scored themselves better) than was objectively measured, whereas under-estimation of performance indicates that the subjects performed worse than was objectively measured. In the case of performance measured as errors committed, a high self-evaluation (i.e., “I committed many errors”), is interpreted as a low score of performance.

### Study quality assessment

To assess the methodological quality on the final set of articles, we used the Quality Assessment Tool for Quantitative Studies (QATQS), which was developed to assess the quality of diverse study designs (Kmet et al., 2004). Each study is scored by the degree to which it satisfies each of the 14 criteria (Yes=2, Partially=1, No=0, Not Applicable=NA). The maximum score an article can receive is calculated as:

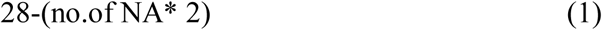

Each article received a summary percent score calculated by dividing sum of the scored items by the maximum possible sum. The assessed items include the adequate reporting of objectives, design, results, and conclusions; the study population and comparator selection methods; the outcomes defined; and statistical methods undertaken. The QATQS does not have a guide for interpreting the final score, therefore we classified study quality as weak (>50%), moderate-weak (51 to 65%), moderate-strong (66 to 79%), or strong (>80%), based on guidelines by de Vet and colleagues (1997), like previous work that used the QATQS (Estabrooks et al., 2009; Squires et al., 2011).

## Results

### Studies reviewed

Twenty-four articles out of the 2849 records yielded from the database searches met the inclusion criteria for this review (Figure 1). The secondary reference list searches did not reveal any additional records. One study consisted of two parts published separately (Bao, Howard, et al., 2006; Bao, Spielholz, et al., 2006), and is treated as a single study for the purposes of this review. Thus, 23 studies were included in the final review. The methodological design of the articles included: 16 cross-sectional designs (70%), three experimental designs (13%), three measurement validity designs (13%), and one descriptive design (4%).

### Study quality

Methodological quality of the included articles is reported in Table 1. Of the 23 included studies, five were rated as strong (22%), 15 as moderate-strong (65%), two as moderate-weak (9%), and one as weak (4%). The included studies were sufficient in describing categories such as study design, subject characteristics, outcome measures and analytic methods, reporting the results and estimate of variance, and stating conclusions that were supported by the results. However, many studies insufficiently described the objective and the following methods details: subject selection and input variables, blinding of investigators when possible, and appropriate sample size. Overall, quality of the included studies was moderate-strong.

**Table 1.**
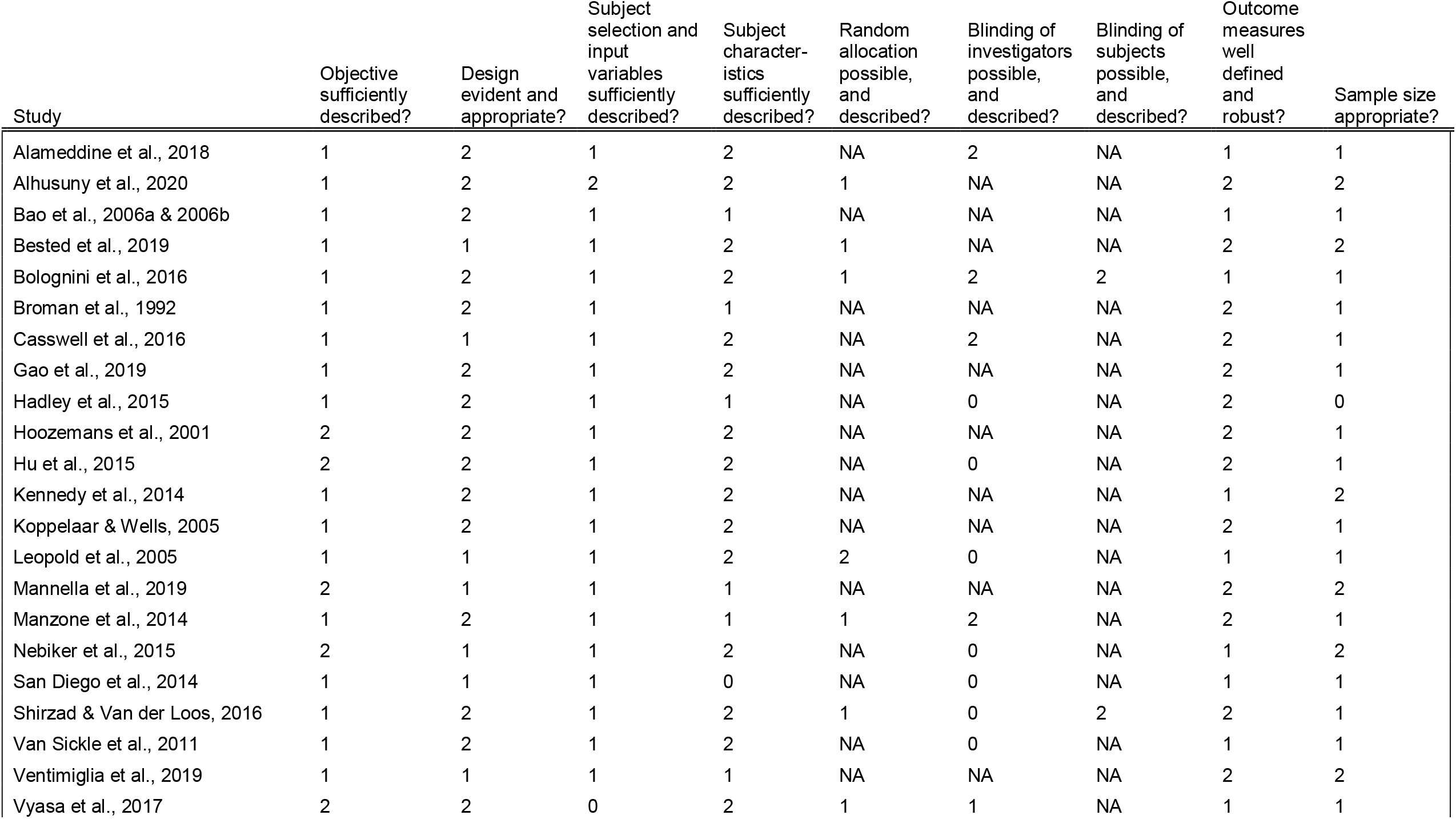
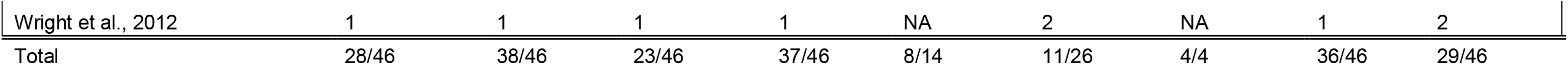
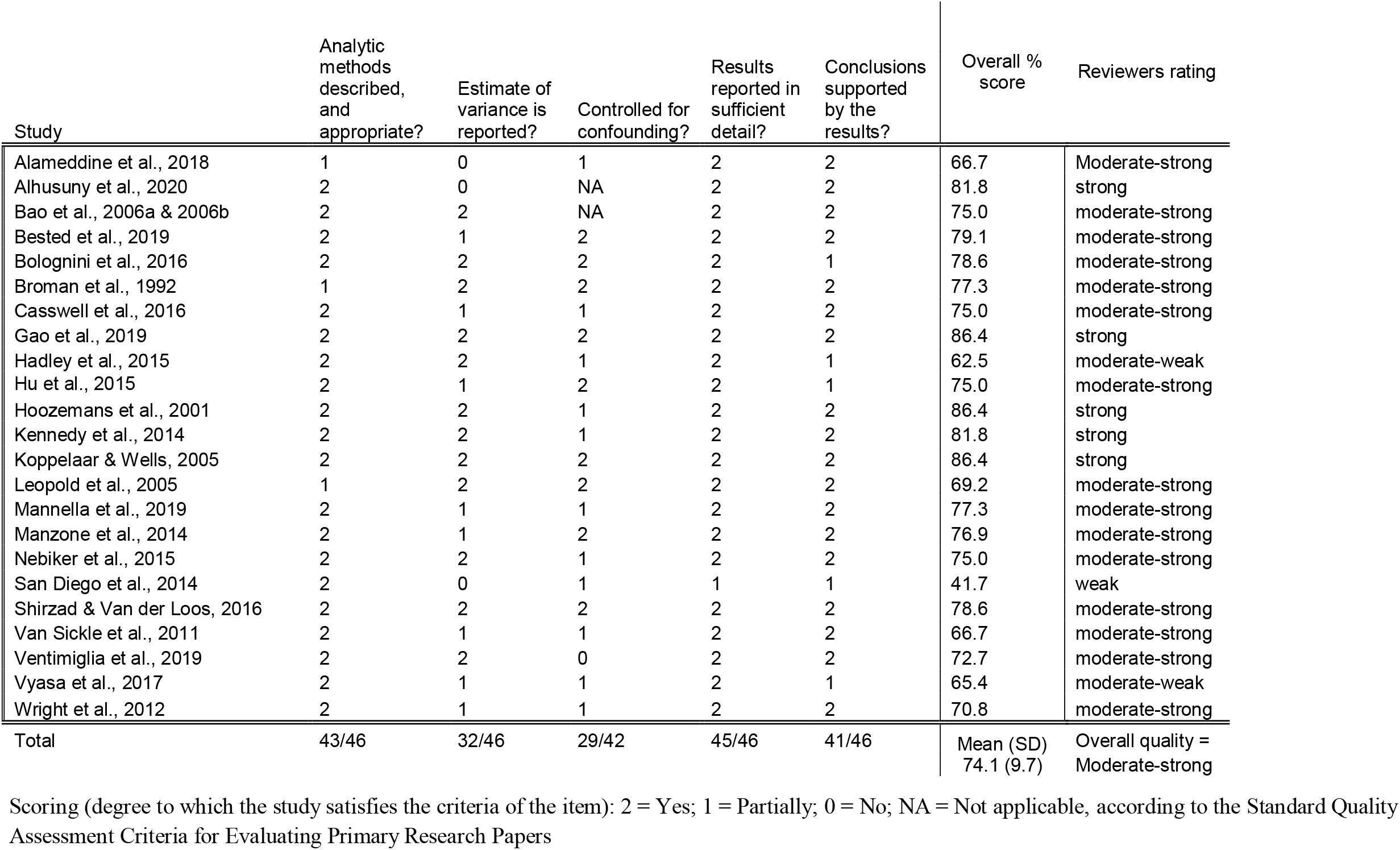
Results of the study quality assessment.

| Study | Objective sufficiently described? | Design evident and appropriate? | Subject selection and input variables sufficiently described? | Subject characteristics sufficiently described? | Random allocation possible, and described? | Blinding of investigators possible, and described? | Blinding of subjects possible, and described? | Outcome measures well defined and robust? | Sample size appropriate? |
| --- | --- | --- | --- | --- | --- | --- | --- | --- | --- |
| Alameddine et al., 2018 | 1 | 2 | 1 | 2 | NA | 2 | NA | 1 | 1 |
| Alhusuny et al., 2020 | 1 | 2 | 2 | 2 | 1 | NA | NA | 2 | 2 |
| Bao et al., 2006a & 2006b | 1 | 2 | 1 | 1 | NA | NA | NA | 1 | 1 |
| Bested et al., 2019 | 1 | 1 | 1 | 2 | 1 | NA | NA | 2 | 2 |
| Bolognini et al., 2016 | 1 | 2 | 1 | 2 | 1 | 2 | 2 | 1 | 1 |
| Broman et al., 1992 | 1 | 2 | 1 | 1 | NA | NA | NA | 2 | 1 |
| Casswell et al., 2016 | 1 | 1 | 1 | 2 | NA | 2 | NA | 2 | 1 |
| Gao et al., 2019 | 1 | 2 | 1 | 2 | NA | NA | NA | 2 | 1 |
| Hadley et al., 2015 | 1 | 2 | 1 | 1 | NA | 0 | NA | 2 | 0 |
| Hoozemans et al., 2001 | 2 | 2 | 1 | 2 | NA | NA | NA | 2 | 1 |
| Hu et al., 2015 | 2 | 2 | 1 | 2 | NA | 0 | NA | 2 | 1 |
| Kennedy et al., 2014 | 1 | 2 | 1 | 2 | NA | NA | NA | 1 | 2 |
| Koppelaar & Wells, 2005 | 1 | 2 | 1 | 2 | NA | NA | NA | 2 | 1 |
| Leopold et al., 2005 | 1 | 1 | 1 | 2 | 2 | 0 | NA | 1 | 1 |
| Mannella et al., 2019 | 2 | 1 | 1 | 1 | NA | NA | NA | 2 | 2 |
| Manzone et al., 2014 | 1 | 2 | 1 | 1 | 1 | 2 | NA | 2 | 1 |
| Nebiker et al., 2015 | 2 | 1 | 1 | 2 | NA | 0 | NA | 1 | 2 |
| San Diego et al., 2014 | 1 | 1 | 1 | 0 | NA | 0 | NA | 1 | 1 |
| Shirzad & Van der Loos, 2016 | 1 | 2 | 1 | 2 | 1 | 0 | 2 | 2 | 1 |
| Van Sickle et al., 2011 | 1 | 2 | 1 | 2 | NA | 0 | NA | 1 | 1 |
| Ventimiglia et al., 2019 | 1 | 1 | 1 | 1 | NA | NA | NA | 2 | 2 |
| Vyasa et al., 2017 | 2 | 2 | 0 | 2 | 1 | 1 | NA | 1 | 1 |
| Wright et al., 2012 | 1 | 1 | 1 | 1 | NA | 2 | NA | 1 | 2 |
| Total | 28/46 | 38/46 | 23/46 | 37/46 | 8/14 | 11/26 | 4/4 | 36/46 | 29/46 |

**Table 1.** Results of the study quality assessment (continued).
| Study | Analytic methods described, and appropriate? | Estimate of variance is reported? | Controlled for confounding? | Results reported in sufficient detail? | Conclusions supported by the results? | Overall % score | Reviewers rating |
| --- | --- | --- | --- | --- | --- | --- | --- |
| Alameddine et al., 2018 | 1 | 0 | 1 | 2 | 2 | 66.7 | Moderate-strong |
| Alhusuny et al., 2020 | 2 | 0 | NA | 2 | 2 | 81.8 | strong |
| Bao et al., 2006a & 2006b | 2 | 2 | NA | 2 | 2 | 75.0 | moderate-strong |
| Bested et al., 2019 | 2 | 1 | 2 | 2 | 2 | 79.1 | moderate-strong |
| Bolognini et al., 2016 | 2 | 2 | 2 | 2 | 1 | 78.6 | moderate-strong |
| Broman et al., 1992 | 1 | 2 | 2 | 2 | 2 | 77.3 | moderate-strong |
| Casswell et al., 2016 | 2 | 1 | 1 | 2 | 2 | 75.0 | moderate-strong |
| Gao et al., 2019 | 2 | 2 | 2 | 2 | 2 | 86.4 | strong |
| Hadley et al., 2015 | 2 | 2 | 1 | 2 | 1 | 62.5 | moderate-weak |
| Hu et al., 2015 | 2 | 1 | 2 | 2 | 1 | 75.0 | moderate-strong |
| Hoozemans et al., 2001 | 2 | 2 | 1 | 2 | 2 | 86.4 | strong |
| Kennedy et al., 2014 | 2 | 2 | 1 | 2 | 2 | 81.8 | strong |
| Koppelaar & Wells, 2005 | 2 | 2 | 2 | 2 | 2 | 86.4 | strong |
| Leopold et al., 2005 | 1 | 2 | 2 | 2 | 2 | 69.2 | moderate-strong |
| Mannella et al., 2019 | 2 | 1 | 1 | 2 | 2 | 77.3 | moderate-strong |
| Manzone et al., 2014 | 2 | 1 | 2 | 2 | 2 | 76.9 | moderate-strong |
| Nebiker et al., 2015 | 2 | 2 | 1 | 2 | 2 | 75.0 | moderate-strong |
| San Diego et al., 2014 | 2 | 0 | 1 | 1 | 1 | 41.7 | weak |
| Shirzad & Van der Loos, 2016 | 2 | 2 | 2 | 2 | 2 | 78.6 | moderate-strong |
| Van Sickle et al., 2011 | 2 | 1 | 1 | 2 | 2 | 66.7 | moderate-strong |
| Ventimiglia et al., 2019 | 2 | 2 | 0 | 2 | 2 | 72.7 | moderate-strong |
| Vyasa et al., 2017 | 2 | 1 | 1 | 2 | 1 | 65.4 | moderate-weak |
| Wright et al., 2012 | 2 | 1 | 1 | 2 | 2 | 70.8 | moderate-strong |
| Total | 43/46 | 32/46 | 29/42 | 45/46 | 41/46 | Mean (SD)<br>74.1 (9.7) | Overall quality =<br>Moderate-strong |

Scoring (degree to which the study satisfies the criteria of the item): 2 = Yes; 1 = Partially; 0 = No; NA = Not applicable, according to the Standard Quality Assessment Criteria for Evaluating Primary Research Papers

### Description of the studies and evaluation tools

#### Upper-extremity task investigated

Table 2 reports the characteristics of interest for each study. Upper extremity tasks that were investigated ranged from simple movements such as finger tapping to complex tasks such as endoscopy.

**Table 2.**
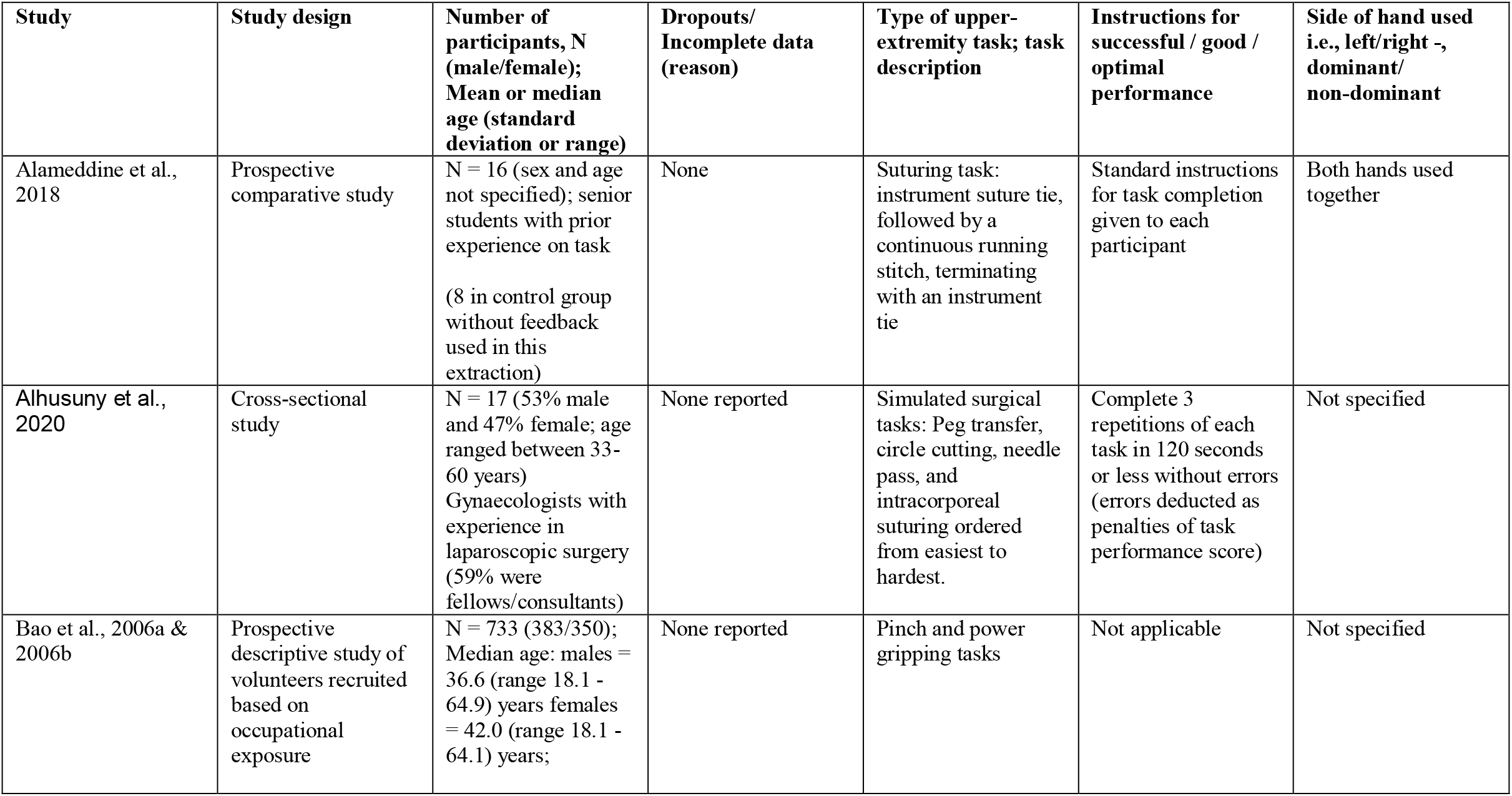
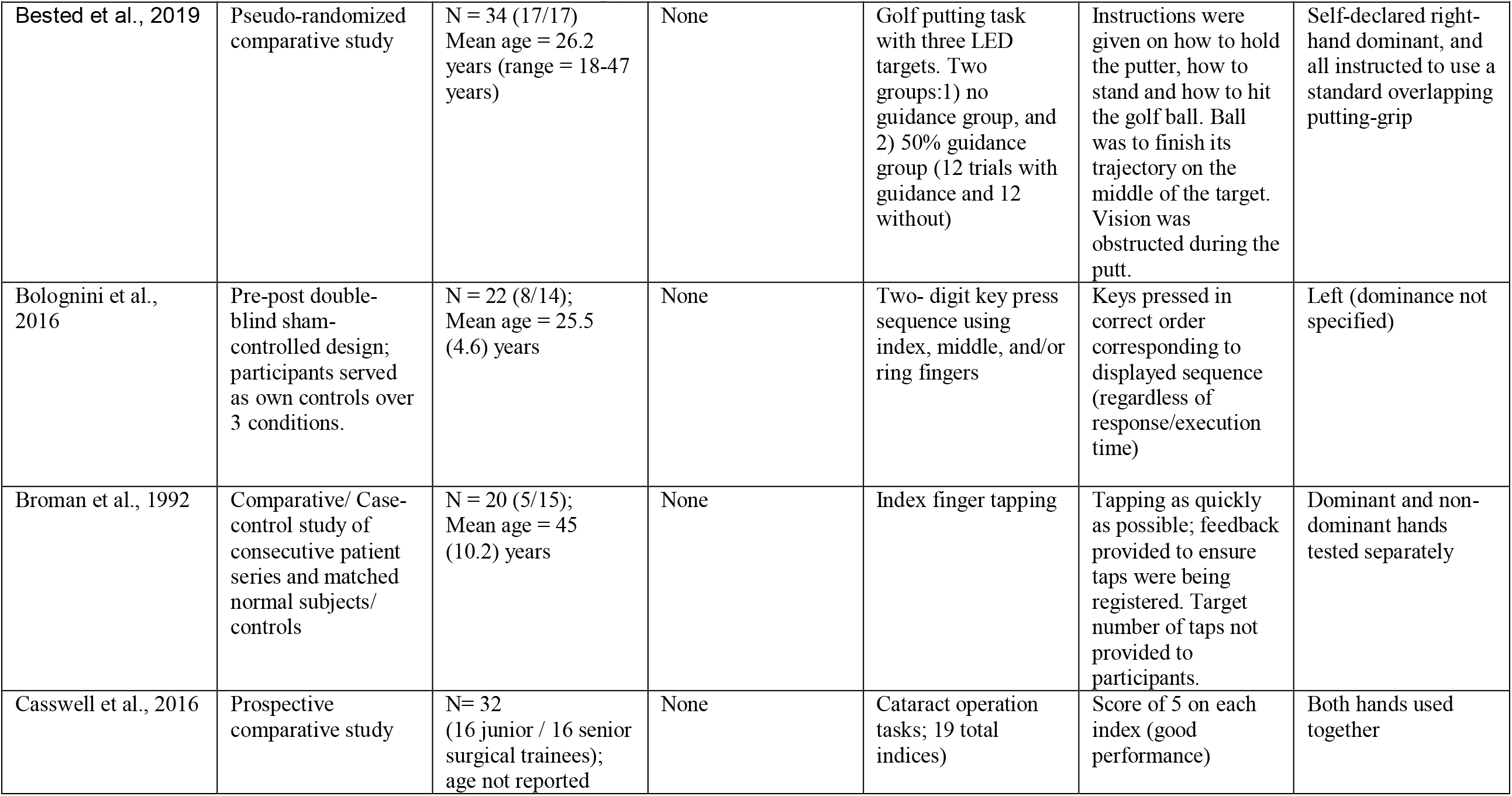
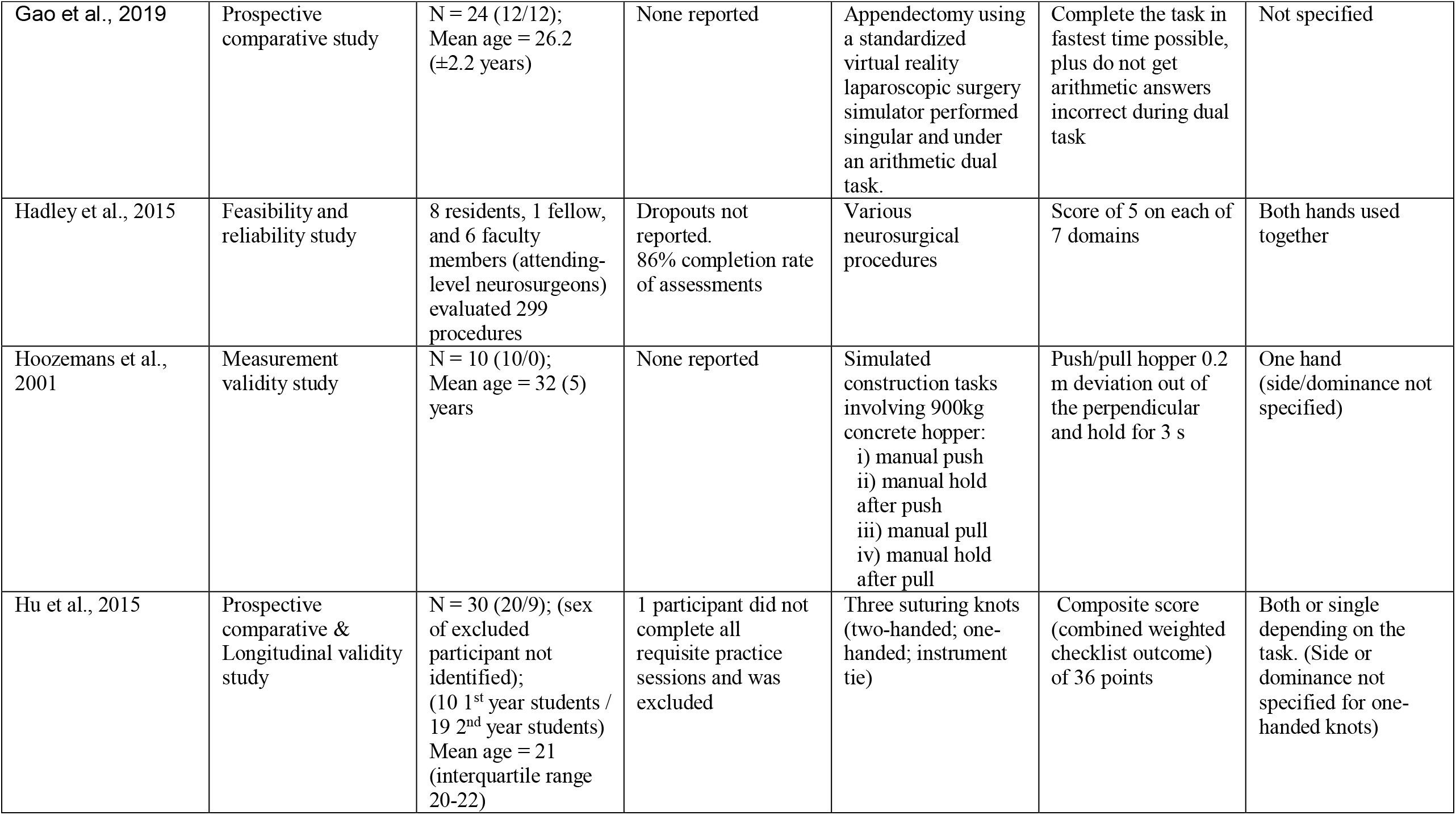
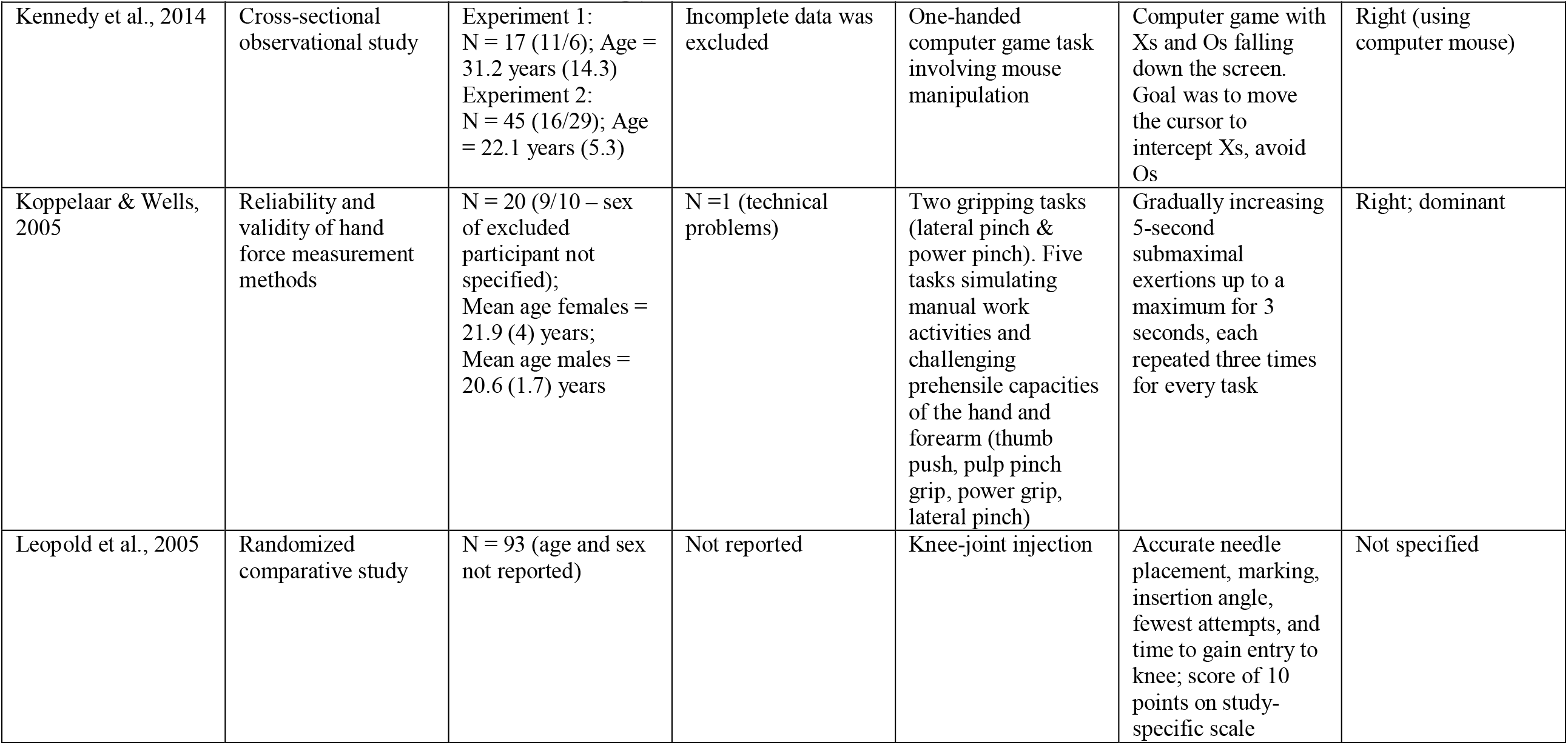
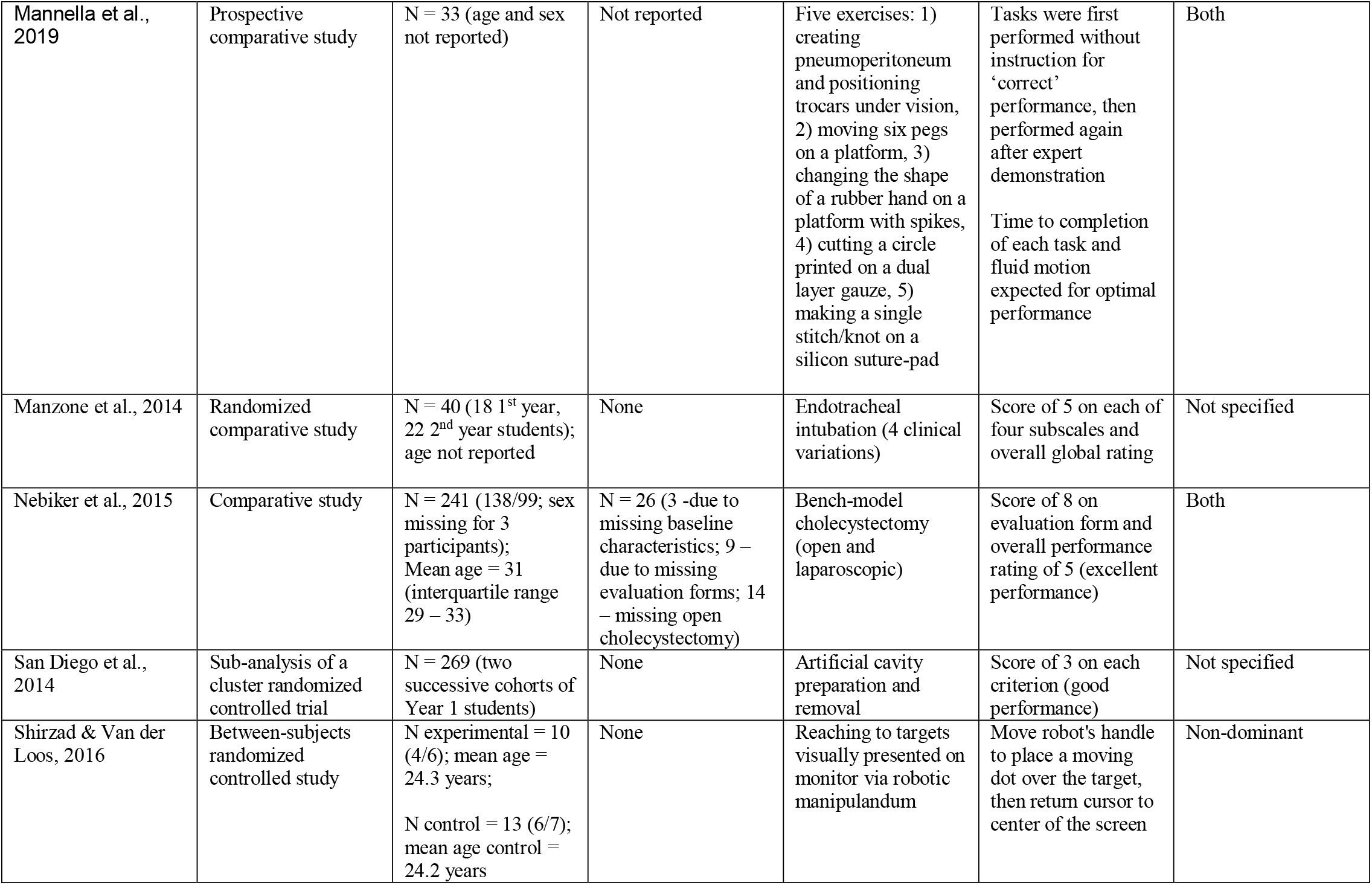
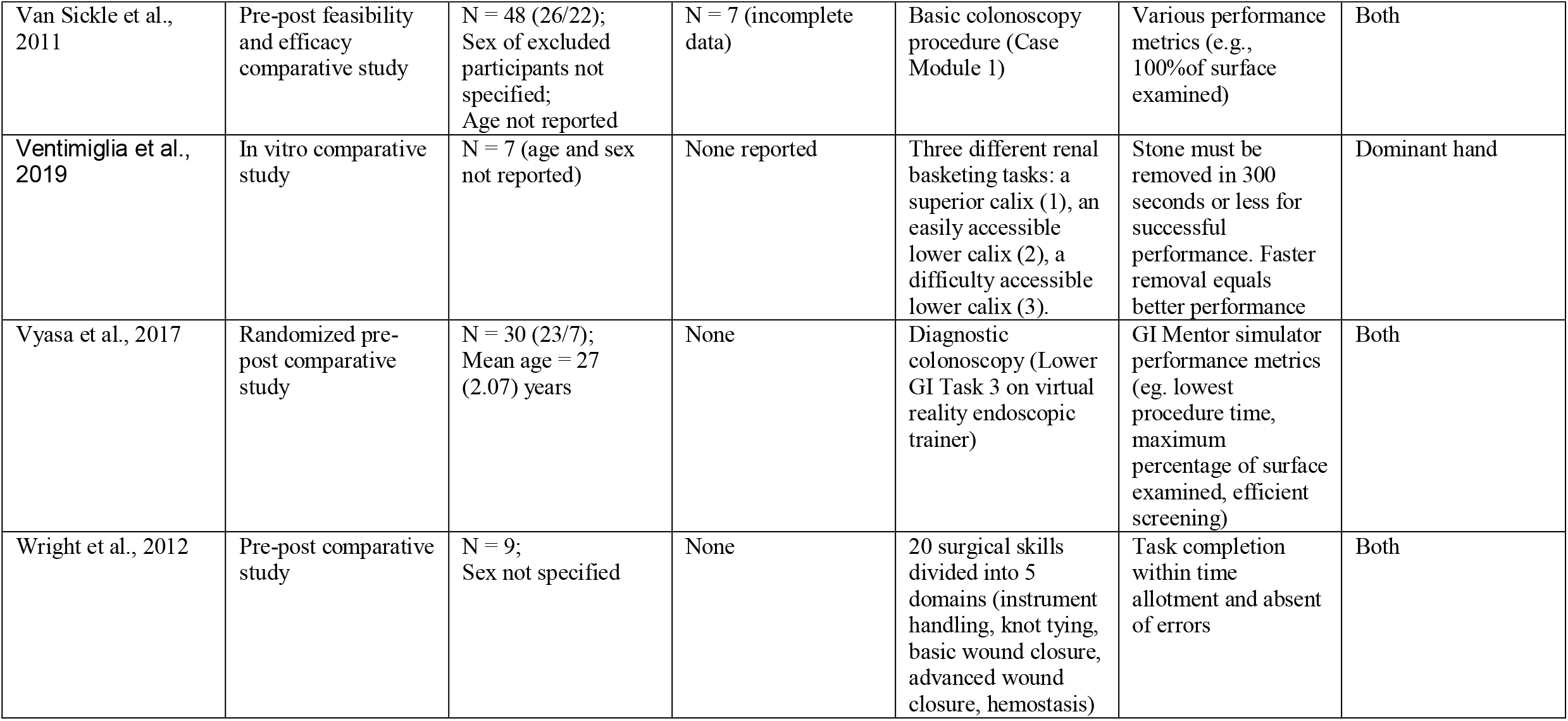
Study characteristics.

| **Study** | **Study design** | **Number of participants, N (male/female); Mean or median age (standard deviation or range)** | **Dropouts/ Incomplete data (reason)** | **Type of upper-extremity task; task description** | **Instructions for successful / good / optimal performance** | **Side of hand used i.e., left/right -, dominant/ non-dominant** |
| --- | --- | --- | --- | --- | --- | --- |
| Alameddine et al., 2018 | Prospective comparative study | N = 16 (sex and age not specified); senior students with prior experience on task<br>(8 in control group without feedback used in this extraction) | None | Suturing task: instrument suture tie, followed by a continuous running stitch, terminating with an instrument tie | Standard instructions for task completion given to each participant | Both hands used together |
| Alhusuny et al., 2020 | Cross-sectional study | N = 17 (53% male and 47% female; age ranged between 33-60 years)<br>Gynaecologists with experience in laparoscopic surgery (59% were fellows/consultants) | None reported | Simulated surgical tasks: Peg transfer, circle cutting, needle pass, and intracorporeal suturing ordered from easiest to hardest. | Complete 3 repetitions of each task in 120 seconds or less without errors (errors deducted as penalties of task performance score) | Not specified |
| Bao et al., 2006a & 2006b | Prospective descriptive study of volunteers recruited based on occupational exposure | N = 733 (383/350); Median age: males = 36.6 (range 18.1 - 64.9) years females = 42.0 (range 18.1 - 64.1) years; | None reported | Pinch and power gripping tasks | Not applicable | Not specified |

| Study | Study design | Number of participants, N (male/female); Mean or median age (standard deviation or range) | Dropouts/ Incomplete data (reason) | Type of upper-extremity task; task description | Instructions for successful / good / optimal performance | Side of hand used i.e., left/right -, dominant/ non-dominant |
| --- | --- | --- | --- | --- | --- | --- |
| Bested et al., 2019 | Pseudo-randomized comparative study | N = 34 (17/17)<br>Mean age = 26.2 years (range = 18-47 years) | None | Golf putting task with three LED targets. Two groups: 1) no guidance group, and 2) 50% guidance group (12 trials with guidance and 12 without) | Instructions were given on how to hold the putter, how to stand and how to hit the golf ball. Ball was to finish its trajectory on the middle of the target. Vision was obstructed during the putt. | Self-declared right-hand dominant, and all instructed to use a standard overlapping putting-grip |
| Bolognini et al., 2016 | Pre-post double-blind sham-controlled design; participants served as own controls over 3 conditions. | N = 22 (8/14);<br>Mean age = 25.5 (4.6) years | None | Two-digit key press sequence using index, middle, and/or ring fingers | Keys pressed in correct order corresponding to displayed sequence (regardless of response/execution time) | Left (dominance not specified) |
| Broman et al., 1992 | Comparative/ Case-control study of consecutive patient series and matched normal subjects/ controls | N = 20 (5/15);<br>Mean age = 45 (10.2) years | None | Index finger tapping | Tapping as quickly as possible; feedback provided to ensure taps were being registered. Target number of taps not provided to participants. | Dominant and non-dominant hands tested separately |
| Casswell et al., 2016 | Prospective comparative study | N= 32 (16 junior / 16 senior surgical trainees); age not reported | None | Cataract operation tasks; 19 total indices) | Score of 5 on each index (good performance) | Both hands used together |
| Gao et al., 2019 | Prospective comparative study | N = 24 (12/12); Mean age = 26.2 ($\pm 2.2$ years) | None reported | Appendectomy using a standardized virtual reality laparoscopic surgery simulator performed singular and under an arithmetic dual task. | Complete the task in fastest time possible, plus do not get arithmetic answers incorrect during dual task | Not specified |
| Hadley et al., 2015 | Feasibility and reliability study | 8 residents, 1 fellow, and 6 faculty members (attending-level neurosurgeons) evaluated 299 procedures | Dropouts not reported. 86% completion rate of assessments | Various neurosurgical procedures | Score of 5 on each of 7 domains | Both hands used together |
| Hoozemans et al., 2001 | Measurement validity study | N = 10 (10/0); Mean age = 32 (5) years | None reported | Simulated construction tasks involving 900kg concrete hopper:<br>i) manual push<br>ii) manual hold after push<br>iii) manual pull<br>iv) manual hold after pull | Push/pull hopper 0.2 m deviation out of the perpendicular and hold for 3 s | One hand (side/dominance not specified) |
| Hu et al., 2015 | Prospective comparative & Longitudinal validity study | N = 30 (20/9); (sex of excluded participant not identified); (10 1<sup>st</sup> year students / 19 2<sup>nd</sup> year students) Mean age = 21 (interquartile range 20-22) | 1 participant did not complete all requisite practice sessions and was excluded | Three suturing knots (two-handed; one-handed; instrument tie) | Composite score (combined weighted checklist outcome) of 36 points | Both or single depending on the task. (Side or dominance not specified for one-handed knots) |
| Kennedy et al., 2014 | Cross-sectional observational study | Experiment 1:<br>N = 17 (11/6); Age = 31.2 years (14.3)<br>Experiment 2:<br>N = 45 (16/29); Age = 22.1 years (5.3) | Incomplete data was excluded | One-handed computer game task involving mouse manipulation | Computer game with Xs and Os falling down the screen. Goal was to move the cursor to intercept Xs, avoid Os | Right (using computer mouse) |
| Koppelaar & Wells, 2005 | Reliability and validity of hand force measurement methods | N = 20 (9/10 – sex of excluded participant not specified);<br>Mean age females = 21.9 (4) years;<br>Mean age males = 20.6 (1.7) years | N =1 (technical problems) | Two gripping tasks (lateral pinch & power pinch). Five tasks simulating manual work activities and challenging prehensile capacities of the hand and forearm (thumb push, pulp pinch grip, power grip, lateral pinch) | Gradually increasing 5-second submaximal exertions up to a maximum for 3 seconds, each repeated three times for every task | Right; dominant |
| Leopold et al., 2005 | Randomized comparative study | N = 93 (age and sex not reported) | Not reported | Knee-joint injection | Accurate needle placement, marking, insertion angle, fewest attempts, and time to gain entry to knee; score of 10 points on study-specific scale | Not specified |

| Study | Study design | Number of participants, N (male/female); Mean or median age (standard deviation or range) | Dropouts/ Incomplete data (reason) | Type of upper-extremity task; task description | Instructions for successful / good / optimal performance | Side of hand used i.e., left/right -, dominant/ non-dominant |
| --- | --- | --- | --- | --- | --- | --- |
| Mannella et al., 2019 | Prospective comparative study | N = 33 (age and sex not reported) | Not reported | Five exercises: 1) creating pneumoperitoneum and positioning trocars under vision, 2) moving six pegs on a platform, 3) changing the shape of a rubber hand on a platform with spikes, 4) cutting a circle printed on a dual layer gauze, 5) making a single stitch/knot on a silicon suture-pad | Tasks were first performed without instruction for 'correct' performance, then performed again after expert demonstration<br>Time to completion of each task and fluid motion expected for optimal performance | Both |
| Manzone et al., 2014 | Randomized comparative study | N = 40 (18 1<sup>st</sup> year, 22 2<sup>nd</sup> year students); age not reported | None | Endotracheal intubation (4 clinical variations) | Score of 5 on each of four subscales and overall global rating | Not specified |
| Nebiker et al., 2015 | Comparative study | N = 241 (138/99; sex missing for 3 participants); Mean age = 31 (interquartile range 29 – 33) | N = 26 (3 -due to missing baseline characteristics; 9 – due to missing evaluation forms; 14 – missing open cholecystectomy) | Bench-model cholecystectomy (open and laparoscopic) | Score of 8 on evaluation form and overall performance rating of 5 (excellent performance) | Both |
| San Diego et al., 2014 | Sub-analysis of a cluster randomized controlled trial | N = 269 (two successive cohorts of Year 1 students) | None | Artificial cavity preparation and removal | Score of 3 on each criterion (good performance) | Not specified |
| Shirzad & Van der Loos, 2016 | Between-subjects randomized controlled study | N experimental = 10 (4/6); mean age = 24.3 years;<br>N control = 13 (6/7); mean age control = 24.2 years | None | Reaching to targets visually presented on monitor via robotic manipulandum | Move robot's handle to place a moving dot over the target, then return cursor to center of the screen | Non-dominant |
| Van Sickle et al., 2011 | Pre-post feasibility and efficacy comparative study | N = 48 (26/22); Sex of excluded participants not specified; Age not reported | N = 7 (incomplete data) | Basic colonoscopy procedure (Case Module 1) | Various performance metrics (e.g., 100% of surface examined) | Both |
| Ventimiglia et al., 2019 | In vitro comparative study | N = 7 (age and sex not reported) | None reported | Three different renal basketing tasks: a superior calix (1), an easily accessible lower calix (2), a difficulty accessible lower calix (3). | Stone must be removed in 300 seconds or less for successful performance. Faster removal equals better performance | Dominant hand |
| Vyasa et al., 2017 | Randomized pre-post comparative study | N = 30 (23/7); Mean age = 27 (2.07) years | None | Diagnostic colonoscopy (Lower GI Task 3 on virtual reality endoscopic trainer) | GI Mentor simulator performance metrics (eg. lowest procedure time, maximum percentage of surface examined, efficient screening) | Both |
| Wright et al., 2012 | Pre-post comparative study | N = 9; Sex not specified | None | 20 surgical skills divided into 5 domains (instrument handling, knot tying, basic wound closure, advanced wound closure, hemostasis) | Task completion within time allotment and absent of errors | Both |

#### Self-evaluation tools used to assess upper-extremity task performance

Table 3 includes data on the measurement tools used to assess performance. The most common self-evaluation tools included Likert-type interval, ratio, or categorical scales (13 studies (Alameddine et al., 2018; Alhusuny et al., 2020; Bao, Howard, et al., 2006; Casswell et al., 2016; Hadley et al., 2015; Leopold et al., 2005; Mannella et al., 2019; Manzone et al., 2014; Nebiker et al., 2015; San Diego et al., 2014; Shirzad & Van der Loos, 2015; Van Sickle et al., 2011; Ventimiglia et al., 2020)). Visual analog scales were used in five studies (Bolognini et al., 2016; Broman et al., 1992; Gao et al., 2019; Kennedy et al., 2014; Koppelaar & Wells, 2005), and another three studies (Bested et al., 2019; Hoozemans et al., 2001; Vyasa et al., 2017) used self-evaluation tools wherein subjects estimated performance numerically (i.e., number of errors made, and amount of force used). Finally, one study used a self-evaluation dichotomous (yes/no) scale (Hu et al., 2015), and one study used a combination of continuous and interval response options in the self-evaluation tool (Wright et al., 2012).

**Table 3.** Study data extraction for the comparison of self- and objective-evaluations of motor performance

| Study | Data Analysis Methodology | Self-Evaluation of Performance |  | Objective Evaluation of Performance |  | Reported comparison between evaluations |
| --- | --- | --- | --- | --- | --- | --- |
|  |  | Outcome Measure | Mean score (SD) | Outcome Measure | Mean Score (SD) |  |
| Alameddine et al., 2018 | Mean pre – post change score (no statistical test performed) | Global Operative Assessment of Laparoscopic Skills (modified for study-specific task) | Overall total: 0.16<br>Bimanual dexterity: 0.25<br>Efficiency: 0.25<br>Tissue handling: 0.13<br>Consistency: 0.00 | Global Operative Assessment of Laparoscopic Skills (modified for study-specific task; by faculty assessor) | Overall total: 0.33<br>Bimanual dexterity: 0.13<br>Efficiency: 0.38<br>Tissue handling: 0.56<br>Consistency: 0.25 | Objective faculty assessors reported greater post-training performance in overall score, plus efficiency, tissue handling, and consistency categories than participant's self-evaluations. Participants self-evaluated post-training performance greater in bimanual dexterity than the faculty assessors. |
| Alhusuny et al., 2020 | No comparison performed | NASA-TLX | Performance subscale not reported | Task performance score = (120 s. × 3 repetitions) – (actual time spent for all three repetitions) – (penalties × 10)<br>(Higher scores indicate better performance). | Total performance<br>2D: 191.8<br>3D: 215.9<br>Peg Transfer<br>2D: 95.6<br>3D: 108.5<br>Accurate Circle Cutting<br>2D: 27.4<br>3D: 29.1<br>Accurate Needle pass<br>2D: 53.5<br>3D: 55.3<br>Intracorporeal Suturing<br>2D: 16.5<br>3D: 22.9 | Not reported |
| Bao et al., 2006a & 2006b | Spearman rank-order correlation between directly measured forces and self-reported force levels | Borg 10-point scale RPE (only pinch and power gripping tasks) | Not reported | Direct measurement (hand dynamometer (N)) | Reported for individual exposure categories – data for correlation analysis not specified | Pinch grip (n=402)<br>Self-report RPE – Directly measured<br>$r = 0.36, p < 0.0001$<br>Power grip (n=322)<br>Self-report RPE – Directly measured<br>$r = 0.36, p < 0.0001$ |
|  |  | Outcome Measure | Mean score (SD) | Outcome Measure | Mean Score (SD) |  |
| Bested et al., 2019 | No comparison performed | Error estimation was assessed by asking the participants to guess where their ball landed. Absolute error estimation in the primary movement axis (AE EstY) pre-test, immediate retention (Imm-Ret), and delay retention (Del-Ret) | No guidance:<br>Pre-test = 69.2 (39) cm<br>Imm-Ret = 49.7 (22) cm<br>Del-Ret = 50.1 (26) cm<br>50% guidance:<br>Pre-test = 58.5 (25) cm<br>Imm-Ret = 40.3 (16) cm<br>Del-Ret = 32.4 (8) cm | A custom grid system was used to determine the exact location where the ball stopped (to the nearest millimeter). Ball endpoint in the primary movement axis (EndY) during the immediate-retention experimental phase. | No guidance:<br>Pre-test = 247 (62) cm<br>Imm-Ret = 247 (26) cm<br>Del-Ret = 255 (36) cm<br>50% guidance:<br>Pre-test = 255 (45) cm<br>Imm-Ret = 230 (21) cm<br>Del-Ret = 216 (23) cm | Evaluations not directly compared. |
| Bolognini et al., 2016 | Pearson's correlation between motor task error rate and self-evaluated performance before and after each tDCS condition | Motor Performance Awareness Questionnaire (100-mm VAS; 2 questions: "I made many errors" and "My performance was fast") | "Errors":<br>Pre-tDCS = 39.9 mm<br>Post-tDCS = 23.7 mm<br>"Fast":<br>Pre-tDCS = 73.3 mm<br>Post-tDCS = 65.92 mm | Error rate;<br>Response time | Error rate:<br>Pre-tDCS = 8% (10%)<br>Post-tDCS = 12% (9%)<br>Response time:<br>Pre-tDCS = 8.39 (1.53) s<br>Post-tDCS = 8.75 (1.67) s | Error Rate:<br>Pre-tDCS<br>Sham, $r = .43$, $p > .05$;<br>PPC, $r = .53$, $p > .05$;<br>PM, $r = .54$, $p > .05$<br>Post-tDCS<br>Sham: $r = .59$, $p = .004$<br>PPC: $r = .69$, $p = .001$<br>PM: $r = .22$, $p = .32$<br>Reaction Time:<br>all p-values $> .11$ (r values not reported) |
| Broman et al., 1992 | No comparison performed | 100-mm VAS; 2 questions: "performance evaluation" | Subjective ratings not specified for subtests | Number of index finger taps | Dominant hand = 61.7 (8.9)<br>Non-dominant hand = 53.1 (7.7) | Not reported |
|  |  | Outcome Measure | Mean score (SD) | Outcome Measure | Mean Score (SD) |  |
| Casswell et al., 2016 | Agreement (Cohen's kappa) between trainee (junior and senior) and assessor evaluations | OSACSS (by trainee) | Not reported | OSACSS (by expert assessor) | Overall median score:<br>63 (junior)<br>72 (senior) (global and task-specific subscores also reported) | (# of indices)<br>Junior (mean kappa = 0.32 [fair])<br>Kappa <0 less than chance (2/19)<br>0-0.2 slight (3/19)<br>0.2-0.4 fair (7/19)<br>0.4-0.6 moderate (4/19)<br>0.6-0.8 substantial (3/19)<br>Senior (mean kappa = 0.45 [moderate])<br>Kappa 0-0.2 slight (1/19)<br>0.2-0.4 fair (7/19)<br>0.4-0.6 moderate (9/19)<br>0.6-0.8 substantial (1/19)<br>0.8-1.0 almost perfect (1/19) |
| Gao et al., 2019 | No comparison performed | NASA-TLX (VAS-style scale; scores range 0-100) | Performance subscale not reported | Performance statistics were recorded by the virtual laparoscopic surgery simulator (calculation and units unclear) | Total performance score:<br>Single tasking (not distracted) = 91.88<br>Dual tasking (distracted) = 83.75 | Not reported. |
| Hadley et al., 2015 | Comparison between resident and assessor evaluations using Wilcoxon signed-rank test to compare resident and faculty evaluations | OSATS (by resident) | Overall score = 3.75 (0.72)<br>Respect for tissue = 3.91 (0.80)<br>Time and motion = 3.60 (0.76)<br>Instrument handling = 3.69 (0.80) | OSATS (by faculty evaluators) | Overall score = 4.04 (0.73)<br>Respect for tissue = 4.15 (0.61)<br>Time and motion = 3.89 (0.76)<br>Instrument handling = 4.06 (0.85) | Overall score – p = 0.036<br>Respect for tissue – p = 0.091<br>Time and motion – p = 0.161<br>Instrument handling – p = 0.068 |
| Hoozemans et al., 2001 | Analysis of variance (F-test) with a repeated measurements design was performed to detect differences between force evaluation methods | Estimation of peak forces in kg (converted to N) | Initial push = 283.7 (114.1) N<br>Hold after push = 377.5 (126.8) N<br>Initial pull = 352.4 (174.0) N<br>Hold after pull = 448.5 (195.3) N | Force transducers (N) | Initial push = 191.0 (35.8) N<br>Hold after push = 266.8 (27.4) N<br>Initial pull = 199.8 (32.9) N<br>Hold after pull = 267.6 (30.6) N | F-test of difference:<br>Initial push: p = 0.029<br>Hold after push: p = 0.019<br>Initial pull: p = 0.023<br>Hold after pull: p = 0.020 |
|  |  | Outcome Measure | Mean score (SD) | Outcome Measure | Mean Score (SD) |  |
| Hu et al., 2015 | Agreement (Cohen's weighted kappa) between self- and trainer or faculty assessments | OSATS (by student) | Not reported | OSATS (by trainer faculty assessor) | Not reported | Self- and trainer assessments: excellent agreement ($\kappa > 0.81$) for composite task score after 22 attempts<br>Self- and faculty assessments: good agreement ($\kappa = 0.74$) for composite score of last attempt<br>Positive difference between self- and trainer-assessments decreases with increasing task attempts ($p = 0.031$) |
| Kennedy et al., 2014 | Repeated measures ANOVAs with linear and quadratic contrasts between JOPs and success rate | JOPs at 7 different speeds levels (Speed 1 slowest) – left to right sliding scale ranging from “none correct” (0%) to “completely correct” (100%) | Experiment 1:<br>JOPs range from 79.11% (17.27) at Speed 1 to 39.33% (23.94) at Speed 7<br>Experiment 2:<br>JOPs range from 84.81% (15.47) at Speed 1 to 35.01% (18.13) at Speed 7 | Hit rate (as measured by the game at each speed) | Experiment 1:<br>Hit Rates range from 95% (5) at Speed 1 to 36% (4) at Speed 7<br>Experiment 2:<br>Hit Rates range from 95% (10) at Speed 1 to 37% (6) at Speed 7 | Experiment 1:<br>$F(1,16) = 64.64, p < .001, \eta^2 = .80$<br>Experiment 2:<br>$F(1,44) = 175.44, p < .001, \eta^2 = .80$ |
| Koppelaar & Wells, 2005 | Pearson correlation between measurement methods | Perception of effort (VAS) | Not reported | Various direct force measurements (N) | Not reported | Direct measurement - Self-report VAS:<br>$r = 0.52$ |
| Leopold et al., 2005 | Not clearly specified | 10-point Likert-style scale, “unsatisfactory” (1) to “very skillful” (10) performance | Not reported | 10-point Likert-style scale with several elements (graded by preceptor) | Pre-instruction<br>Experienced 5.81 (2.58)<br>Novice 6.59 (2.81)<br>Post-instruction<br>Experienced 8.78 (1.66)<br>Novice 7.85 (2.63) | Not reported |
|  |  | Outcome Measure | Mean score (SD) | Outcome Measure | Mean Score (SD) |  |
| Mannella et al., 2019 | Evaluation scores tested for differences but unclear what statistical procedures were used | OSATS (by resident/trainee) | Exercise 1:<br>Before expert demonstration (A) = 8.303<br>After expert demonstration (B) = 10.06<br>Exercise 2:<br>A = 10.58<br>B = 12.73<br>Exercise 3:<br>A = 11.06<br>B = 13.42<br>Exercise 4:<br>A = 9.485<br>B = 11.88<br>Exercise 5:<br>A = 7.303<br>B = 9.34 | OSATS (by external observer) | Exercise 1:<br>A = 8.697<br>B = 9.03<br>Exercise 2:<br>A = 10.94<br>B = 13.48<br>Exercise 3:<br>A = 11.42<br>B = 14.27<br>Exercise 4:<br>A = 9.606<br>B = 12.06<br>Exercise 5:<br>A = 10.79<br>B = 14.67 | Self-evaluation was significantly lower than objective evaluation scores on exercise 5 (making a single stitch/knot on a silicon suture-pad) only.<br>Exercise 1:<br>A: p = 0.9921<br>B: p = 0.8234<br>Exercise 2:<br>A: p = 0.9921<br>B: p = 0.9176<br>Exercise 3:<br>A: p = 0.9921<br>B: p = 0.9050<br>Exercise 4:<br>A: p = 0.9921<br>B: p = 0.9921<br>Exercise 5:<br>A: p = 0.0001<br>B: p < 0.0001 |
| Manzone et al., 2014 | No comparison performed | Post-study questionnaire | Not reported | Hand motion; global rating scale | Not reported | Not reported |
| Nebiker et al., 2015 | Agreement (Cohen's kappa) between trainee and expert tutor assessments | 5-point overall performance rating scale (by trainee) | Not reported | Objective structured evaluation form (8-item task-specific checklist and 5-point overall performance rating by tutor) | Not specified. | Laparoscopic task<br>Raw agreement: 50% (95% CI: 44%, 57%)<br>Kappa: 0.384 [fair] (95% CI: 0.304, 0.464)<br>Open surgery task<br>Raw agreement: 57% (95% CI: 50%, 64%)<br>Kappa: 0.340 (95% CI: 0.256, 0.423) |
| San Diego et al., 2014 | Agreement (Kw) between staff and student ratings | Evaluation checklist (3-point scale for 5 criteria) by students | Not reported | Evaluation checklist (3-point scale for 5 criteria) by staff assessors) | Not reported | Evaluation criteria: % agreement; Kw<br>Hand-held tooth task (n=138 self under/over-estimate)<br>Angle of entry (67.7%; 0.08; under)<br>Holding instrument (67.4%; 0.19; over)<br>Removal of caries, wall (86.2%; 0.16; over)<br>Removal of caries, floor (92.8%; 0.25; over)<br>Avoid pulp exposure (97.1%; 0.70; agree) |
|  |  | Outcome Measure | Mean score (SD) | Outcome Measure | Mean Score (SD) |  |
|  |  |  |  |  |  | Tooth in-jaw task (n=131)<br>Angle of entry (47.9%; 0.01; under)<br>Holding instrument (37.7%; 0.1; over)<br>Removal of caries, wall (80.5%; 0.48; over)<br>Removal of caries, floor (89.4%; 0.28; over)<br>Avoid pulp exposure (87.8%; 0.16; under) |
| Shirzad & Van der Loos, 2016 | No comparison performed | NASA-TLX – “task performance” subscale (Likert–style scale) | Control group: 2.54 (0.96)<br>Experimental group: 3.40 (0.97) | Reaching accuracy (average maximum deviation) | Not reported | Not available |
| Van Sickle et al., 2011 | No comparison performed | Self-assessment questionnaire (0-4 Likert style-scale) | Pre-training 1.5 (1.0)<br>Post-training 2.7 (0.6) | Total errors measured by GI Mentor II; Global Assessment of Gastrointestinal Endoscopic Skills (5-point Likert-style scale checklist) | Pre-training = 14.9 (2.4)<br>Post-training = 19.6 (1.5) | Not reported |
| Ventimiglia et al., 2019 | Mann-Whitney test and Chi-square test, followed by a subgroup analysis. | NASA-TLX - “task performance” subscale (Likert–style scale; score 1-20) | Assistant basketing:<br>Resident = 13<br>Urologist = 14<br>Overall = 14<br>Lithovue empower:<br>Resident = 14<br>Urologist = 19<br>Overall = 14 | Time from ureteroscope insertion in the K-Box up until stone removal from the K-Box (stone retrieval time in seconds). | Assistant basketing:<br>Task 1 = 60 (27-158)<br>Task 2 = 68 (38-128)<br>Task 3 = 189 (70-247)<br>Lithovue empower:<br>Task 1 = 48 (39-102)<br>Task 2 = 55 (34-81)<br>Task 3 = 123 (60-233) | Not reported |
| Vyasa et al., 2017 | Discrepancy scores (self-assessment score – objective score) | Self-assessment open response questionnaire (1 item for each parameter measured by GI Mentor II system) | Not reported | GI Mentor II simulator computer system measured parameters | All subjects, post-training:<br>Overall time (min)= 15.1 (5.1)<br>Time to cecum (min) = 11.2 (4.9)<br>Screening efficiency (%) = 62.0 (23.9) | Discrepancy scores (all subjects, post-training):<br>Overall time (s) = 44.3 (196.1)<br>Time to cecum (s) = 38.1 (138.8)<br>Screening efficiency (%) = 8.9 (28.0)<br>Surface examined (%) = -14.8 (16.7)<br>Patient in pain time (%) = 14.6 (15.9)<br>On average, subjects overestimated performance in overall completion time, time to cecum, efficiency, and % of patient in pain time. Subjects |
|  |  | Outcome Measure | Mean score (SD) | Outcome Measure | Mean Score (SD) |  |
|  |  |  |  |  | Surface examined (%) = 91.0 (5.5)<br>Patient in pain time (%) = 2.7 (6.2) | underestimated performance on % mucosal surface examined. |
| Wright et al., 2012 | No comparison performed | Author-developed self-assessment tool based on defined time and error metrics (task score = time allowed – [time elapsed + # of errors] x 20); overall score = of 5 task scores | Overall score:<br>Pre-test = 3754 (1742)<br>Post-test = 6496 (1337) | Expert assessment tool (task-specific checklist and anchored global rating scale) | Overall score:<br>Pre-test = 10.1 (2.6)<br>Post-test = 14.6 (2.7) | None/Not available |

Abbreviations: RPE = rating of perceived exertion; N = newtons; VAS = visual analog scale; tDCS = transcranial direct current stimulation; OSACSS = Objective structured assessment of cataract surgical skill; OSATS = Objective structured assessment of technical skill; JOP = judgment of performance; CI = confidence interval; Kw = Cohen's weighted-kappa statistic; NASA-TLX = National Aeronautics and Space Administration – Task Load Index.

Table 4 reports a summary of the number of items regarding performance and the number of response options available to the participant to evaluate their performance for that item. Performance evaluation ranged from a single item or question, such as (paraphrasing) “how did you perform?” or “how much force did you use to perform this action?” (13 studies (Alameddine et al., 2018; Alhusuny et al., 2020; Bao, Howard, et al., 2006; Bested et al., 2019; Gao et al., 2019; Hoozemans et al., 2001; Koppelaar & Wells, 2005; Leopold et al., 2005; Manzone et al., 2014; Nebiker et al., 2015; Shirzad & Van der Loos, 2015; Van Sickle et al., 2011; Ventimiglia et al., 2020)) to nineteen (Casswell et al., 2016) or twenty (Hu et al., 2015) items, wherein the overall performance was broken down into several components. There was also variability in how precisely participants graded themselves.

**Table 4.** Comparison of the self- and objective-evaluation measurements.

| **Study** | **Number of items about task performance** | **Response scale used (continuous, dichotomous, etc.)** | **Number of response options per item** | **Does the self-evaluation methods match objective scoring?** | **Does the self-evaluation measure assess same construct as the objective measure?** | **If items/ responses were missing, was that discussed?** | **Did subjects have knowledge of their objective performance results?** |
| --- | --- | --- | --- | --- | --- | --- | --- |
| Alameddine et al., 2018 | 4 | interval | 5 | Yes | Yes | No missing responses reported | No (control group) |
| Alhusuny et al., 2020 | 1 | interval | 20 | No | No | No missing responses reported | No |
| Bao et al., 2006a & 2006b | 1 | ratio | 10 | No | No | No missing responses reported | Not reported |
| Bested et al., 2019 | 1 | continuous | infinite response options | Yes | Yes | Yes, one participant was removed due to recording errors | No |
| Bolognini et al., 2016 | 2 | VAS (100mm) | 100 | No | No | No missing responses reported | No |
| Broman et al., 1992 | 4 | VAS (100mm) | 100 | No | No | No missing responses reported | Yes (taps registered on computer screen) |
| Casswell et al., 2016 | 19 | interval | 5 | Yes | Yes | No missing responses reported | No |
| Gao et al., 2019 | 1 | VAS (100mm) | 100 | No | No | No missing responses reported | Unclear if feedback provided during testing trials |
| Hadley et al., 2015 | 7 | interval | 5 | Yes | Yes | No missing responses reported | Unclear if feedback provided during practice would influence self-evaluation |
| Hoozemans et al., 2001 | 1 | continuous | infinite response options | No | Yes | No missing responses reported | Not reported |
| Hu et al., 2015 | 20 | dichotomous | YES/NO - point(s) for successful completion of task item | Yes | Yes | No missing responses reported | No |
| Kennedy et al., 2014 | 1 | VAS | Cannot be determined (sliding scale from 'none correct' to 'completely correct') | No | Yes | Yes, some missing data were removed from analysis | No |
| Koppelaar & Wells, 2005 | 1 | VAS (100mm) | 100 | No | No | No missing responses reported | Not reported |
| Leopold et al., 2005 | 1 | interval | 10 | No | No | No missing responses reported | Unclear if feedback provided during testing trials |
| Mannella et al., 2019 | 3 - 5 | interval | 5 | Yes | Yes | No missing responses reported | No |
| Manzone et al., 2014 | 1 | interval | 5 | No | No | No missing responses reported | Unclear if feedback provided during testing trials |
| Nebiker et al., 2015 | 1 | interval | 5 | Yes | Yes | Yes data missing, but unclear how it has handled | No |
| San Diego et al., 2014 | 5 | categorical | 3 (Poor/Adequate/Good) | Yes | Yes | No missing responses reported | Not reported |
| Shirzad & Van der Loos, 2016 | 1 | interval | 5 | No | No | No missing responses reported | Unclear if feedback provided during testing trials would influence self-evaluation |
| Van Sickle et al., 2011 | 1 | ratio | 5 | No | No | No missing responses reported | Not reported |
| Ventimiglia et al., 2019 | 1 | interval | 20 | No | No | No missing responses | Unclear if feedback provided during testing trials |
| Vyasa et al., 2017 | 7 (5 used for analysis) | Continuous / interval | 2 items infinite (time), 3 items % out of 100 / 2 items 10 response options (not used in comparison) | Yes (scores for questions on tool match objective scoring) | Yes | No missing responses reported | No |
| Wright et al., 2012 | 2 (total task time and total task errors) | Other (predefined scoring system based on perceived time elapsed and task errors) | Infinite response options | No | No | No missing responses reported | No |

Abbreviations: VAS = visual analog scale.

#### Objective evaluation of upper-extremity task performance

Objective evaluation was commonly computed by a device or equipment used in the upper extremity motor task of interest. Performance directly scored by the device (e.g., GI Mentor II) was used in 9/23 studies (39%) (Alhusuny et al., 2020; Bolognini et al., 2016; Broman et al., 1992; Gao et al., 2019; Kennedy et al., 2014; Shirzad & Van der Loos, 2015; Van Sickle et al., 2011; Ventimiglia et al., 2020; Vyasa et al., 2017) whereas force production was measured in three studies (13%) (Bao, Howard, et al., 2006; Bao, Spielholz, et al., 2006; Hoozemans et al., 2001; Koppelaar & Wells, 2005) using a force transducer. The remainder of the studies (10/23, 43%) measured task performance with Likert-type objective evaluation tools scored by expert assessors/trainers (Alameddine et al., 2018; Casswell et al., 2016; Hadley et al., 2015; Hu et al., 2015; Leopold et al., 2005; Mannella et al., 2019; Manzone et al., 2014; Nebiker et al., 2015; San Diego et al., 2014; Wright et al., 2012).

#### Concordance between self- and objective-evaluation tools

Table 4 summarizes the similarities and differences between the self- and objective evaluation tools used. Also summarized is the performance construct which refers to the dimension by which performance was measured (e.g., amount of force used, time to task completion, number of performance errors committed).

In eight studies (Alameddine et al., 2018; Bested et al., 2019; Casswell et al., 2016; Hadley et al., 2015; Hu et al., 2015; Mannella et al., 2019; Nebiker et al., 2015; San Diego et al., 2014), performance was self- and objectively evaluated using the same tool (e.g., checklist with Likert-style questions such as the objective structured assessment of technical skill; OSATS). The majority (nine studies (Bao, Spielholz, et al., 2006; Broman et al., 1992; Koppelaar & Wells, 2005; Leopold et al., 2005; Manzone et al., 2014; Shirzad & Van Der Loos, 2012; Van Sickle et al., 2011; Wright et al., 2012)) scored self- and objective-evaluation of performance differently. For instance, Boa et al (2006) measured grip force performance objectively using force transducers, but participants self-evaluated the force level of the grip forces using a rating of perceived exertion on a scale of 1 – 10.

#### Descriptive summary of study findings

Table 5 provides a descriptive summary of the seven studies that contained sufficient information to comment on the accuracy of self-evaluated performance (whether the study subjects self-evaluated their performance lower, higher, or in agreement with the objective evaluation). Four of the studies (Hoozemans et al., 2001; Hu et al., 2015; San Diego et al., 2014; Vyasa et al., 2017) reported on average across participants an overestimation of performance compared to the objective evaluation. Three studies (Hadley et al., 2015; Kennedy et al., 2014; Nebiker et al., 2015) reported an underestimation of performance. In general, there is no clear trend across studies for consistent under- or overestimation of performance compared to an objective measure of the motor task.

**Table 5.**
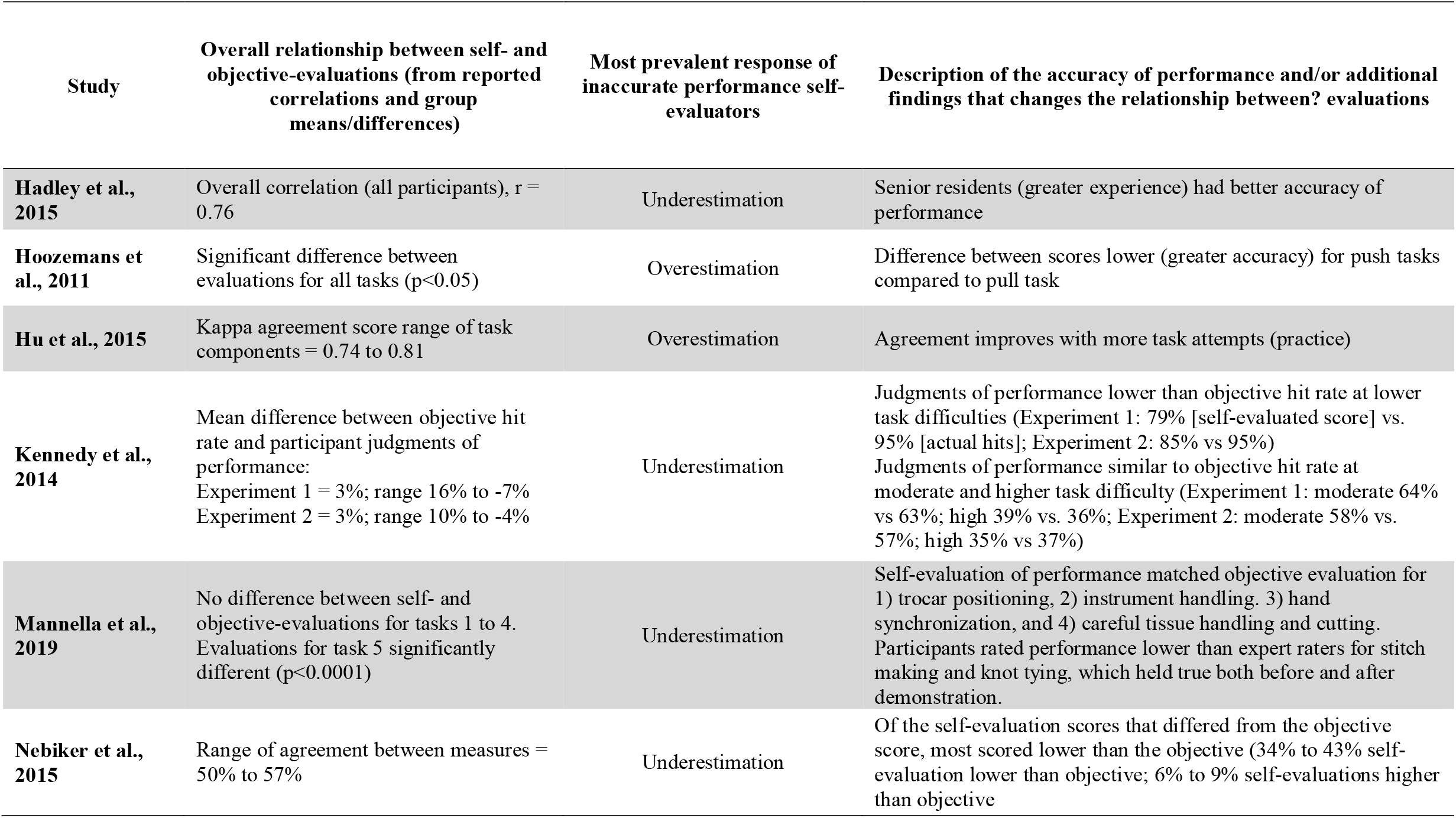
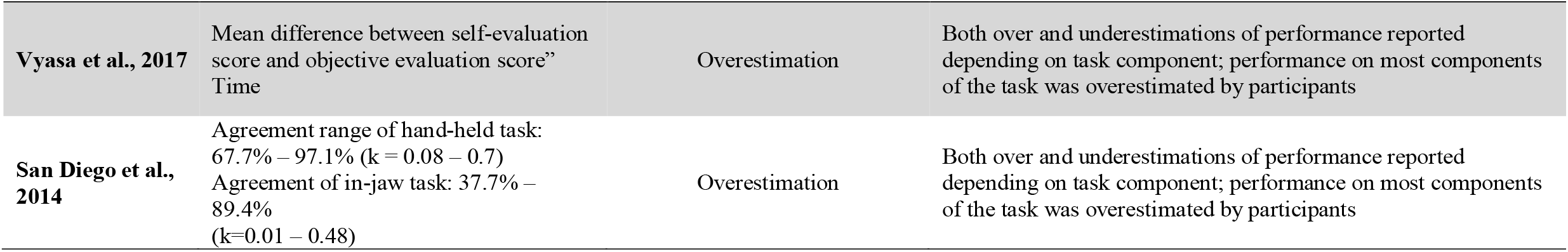
Over- or underestimation of self-evaluation compared to objective evaluation

#### Statistical comparisons of self-evaluation and objective evaluation

Eleven studies reported a statistical comparison between self- and objective evaluations that could be used in this analysis (Bao, Howard, et al., 2006; Bolognini et al., 2016; Casswell et al., 2016; Hadley et al., 2015; Hoozemans et al., 2001; Hu et al., 2015; Kennedy et al., 2014; Koppelaar & Wells, 2005; Nebiker et al., 2015; San Diego et al., 2014; Vyasa et al., 2017) (Table 3). Studies reported positive correlation coefficients ranging from 0.36-0.76 (Bao, Howard, et al., 2006; Bolognini et al., 2016; Hadley et al., 2015; Koppelaar & Wells, 2005) indicating a low to high strength of association between evaluations (Mukaka, 2012). Mean kappa agreements between evaluations ranged from 0.21-0.50, indicating fair to moderate agreement (McHugh, 2012). Three studies reported statistically significant differences between values for self- and objective measures (Hadley et al., 2015; Hoozemans et al., 2001; Vyasa et al., 2017). Finally, Alameddine and coworkers (2018) examined how self- and objective evaluations changed over a period of training. They found a mean objective pre-to-post training performance improvement of 0.33 points, whereas participants’ overall self-evaluated performance improvement was 0.16 points.

#### Over- or underestimation of self-evaluation compared to objective evaluation

Eight studies (Hadley et al., 2015; Hoozemans et al., 2001; Hu et al., 2015; Kennedy et al., 2014; Mannella et al., 2019; Nebiker et al., 2015; San Diego et al., 2014; Vyasa et al., 2017) provided sufficient information to comment on whether the subjects self-evaluated their performance lower (underestimation of performance), higher (overestimation), or in perfect agreement with the objective evaluation (Table 5). Two studies reported overestimations of performance. Hoozemans et al. (Hoozemans et al., 2001) found, while still an overestimation, self-evaluation of force production was more accurate for a pushing task compared to a pulling task. Hu and colleagues (Hu et al., 015) reported large overestimations of self-evaluations during early attempts at a suturing task, with accuracy improving with more task attempts. In contrast, two studies reported underestimation of performance on certain neurosurgical tasks (Hadley et al., 2015; Mannella et al., 2019). While one study noted better self-evaluation accuracy for senior residents compared to juniors (Hadley et al., 2015), the other observed no change in agreement between evaluations, comparing before and after demonstration (Mannella et al., 2019). Two studies found over- and underestimations of performance, depending on the component of the task that was analyzed, but both reported an overestimation of performance on most task components (San Diego et al., 2014; Vyasa et al., 2017). On a computer mouse manipulation task, Kennedy and coworkers (Kennedy et al., 2014) found fairly accurate judgments of performance overall, with most inaccurate self-evaluators underestimating their performance, especially on lower difficulty tasks. The final study described that 50% and 57% of the subjects had perfect agreement with objective evaluation on laparoscopic and open surgery tasks, respectively. However, individuals whose self-evaluation was not in agreement with the objective expert rater (43% of participants in the laparoscopic task and 34% in the open surgery task), significantly underestimated their performance (Nebiker et al., 2015). In summary, of the eight studies included in this sub analysis, four reported an underestimation of performance and four reported an overestimation of performance as the most prevalent kind of inaccurate self-evaluation.

#### Potential factors that influence self-evaluation

Given the heterogeneity of studies included in a scoping review, results charting may be further refined toward the end of the review when the reviewers have the greatest awareness of the contents of the included studies (Peters et al., 2015). Upon completion of data extraction, none of the included studies were conducted with the objective of investigating factors that influence the accuracy of self-evaluation of motor performance. Therefore, the secondary objective to determine factors that may influence motor performance self-evaluation could not be directly assessed based on the conclusions of the included studies. However, one investigator (LC) examined study elements such as participant characteristics, study design, and how the task was performed, to qualitatively identify potential influencing factors based on interpretation of reported results. For example, if a study reported a difference in the accuracy of self-evaluation between two study groups categorized as novice and expert, then experience with the task was identified as a potential factor that influences self-evaluation accuracy. Identified factors include degree of task difficulty (Kennedy et al., 2014), task demonstration (Bested et al., 2019; Mannella et al., 2019), experience with the task (Casswell et al., 2016; Hadley et al., 2015; Hu et al., 2015; Mannella et al., 2019; Manzone et al., 2014; Ventimiglia et al., 2020), relationship of the motor task to occupation or negative consequence of unsuccessful performance (studies involving medical/surgical tasks), and differences in the way subjective and objective performance is measured.

## Discussion

The main objective of this review was to investigate the nature and accuracy of self-evaluation of upper-extremity motor performance by exploring studies that have measured both objective motor performance and self-evaluation of a motor task performed by healthy adults. Most studies included in this review (20 of 23) were of moderate-strong quality or better. Across the 23 studies we observed an association between self- and objective evaluation, however the strength of the correlations varied from weak to strong, agreement scores varied from fair to moderate, and significant differences were found between group means for self- and objective evaluations.

Furthermore, there was a considerable range in correlation coefficients for the relationship between self- and objective-evaluation scores across the studies (r = 0.36 to 0.76; k = 0.21 to 0.50). It is evident that self-evaluation of upper extremity motor performance is not always precisely aligned with the objective measurement of that performance and can be described as moderately accurate. It is important to note that of the included studies, only one (San Diego et al., 2014) had the primary purpose of comparing an individual’s perception of their task performance to an objective measure of that performance. All other studies either explored this question as a secondary objective, or briefly discussed the relationship with little detail. Therefore, this research question has yet to be fully explored. Our review revealed several methodological considerations for further investigations of this relationship. The secondary aim of this review was to describe factors that potentially influence motor performance evaluation. Factors identified from the included studies are task purpose, familiarity, difficulty, and demonstration.

### Methodological considerations

Eleven studies measured both self- and objective-evaluation of performance using the same motor performance criteria or tool (Alameddine et al., 2018; Bested et al., 2019; Casswell et al., 2016; Hadley et al., 2015; Hoozemans et al., 2001; Hu et al., 2015; Kennedy et al., 2014; Mannella et al., 2019; Nebiker et al., 2015; San Diego et al., 2014; Vyasa et al., 2017). Using the same or reliably comparable measurement tools in this way allowed a direct comparison between self- and objective-evaluation methods while maintaining high internal consistency (the degree to which test items measure the same construct) between methods (Henson, 2001). For some of the tasks, it was not possible to assess both self- and objective-evaluation of task performance on the same measurement tool. For example, when performance was measured objectively with a force transducer, subjective ratings were quantified by asking the participant to estimate the amount of force they used in Newtons (Bao, Howard, et al., 2006; Hoozemans et al., 2001; Koppelaar & Wells, 2005). This type of estimate seems somewhat removed from how individuals may think of their motor performance in daily life. This difference in self- and objective-evaluation methods and units of measure may influence the accuracy of self-evaluation.

The remaining twelve (Alhusuny et al., 2020; Bao, Spielholz, et al., 2006; Bolognini et al., 2016; Broman et al., 1992; Gao et al., 2019; Koppelaar & Wells, 2005; Leopold et al., 2005; Manzone et al., 2014; Shirzad & Van der Loos, 2015; Van Sickle et al., 2011; Ventimiglia et al., 2020; Wright et al., 2012) studies measured self- and objective-evaluation using different tools that assessed different aspects or domains of performance. For example, some studies asked participants general questions about performance (e.g., “How well did you perform?” from the NASA-TLX performance domain), while the objective measurement quantified more specific domains of performance (e.g., error in movement trajectory, direct force output, or time to completion minus error penalties) (Alhusuny et al., 2020; Koppelaar & Wells, 2005; Shirzad & Van der Loos, 2015). A more open question about general performance may have the participants considering aspects of their performance that are not related to the construct assessed in the objective measure, which may account for the differences observed between the self- and objective-evaluations of performance. Moreover, a person may self-evaluate their performance using the criteria/units that are of importance to them and their motor abilities, rather than the measure/units the researchers selected for their objective measure. These methodological issues complicate the interpretation of the nature and accuracy of self-evaluation of motor performance.

### Familiarity with the task

Six of the fifteen studies that used tasks specific to a profession, involved participants at varying degrees of experience with the task (i.e., senior versus junior trainees) (Casswell et al., 2016; Hadley et al., 2015; Hu et al., 2015; Mannella et al., 2019; Manzone et al., 2014; Ventimiglia et al., 2020). In general, self-evaluation by senior trainees had better agreement with objective evaluations than junior trainees. This makes sense from a motor learning perspective as greater experience with the task enables great error detection and overall skill mastery (Schmidt, 2003). One study (Ventimiglia et al., 2020) reported no differences in self-evaluation between training levels, despite better objective measures for senior vs junior trainees. It is unclear, however, whether optimal performance was adequately explained to the participants of this study (Ventimiglia et al., 2020), which may have affected their judgements. Ultimately, the relationship between experience and accuracy of self-evaluation of performance may be related to the cumulative amount of practice.

Hu et al. (Hu et al., 2015) aimed to examine the longitudinal development of technical self-assessment by comparing objective- and self-evaluations throughout suturing task training. They found that the difference between self- and objective-evaluations decreases with task practice (p=0.031) with agreement scores reaching good to excellent agreement after 18 to 22 task attempts (Hu et al., 2015). Practice improves performance (Schmidt & Lee, 2011), which may be a result of increased awareness of motor performance. As a performer becomes more experienced with a task, they become better at identifying performance errors (Schmidt & Lee, 2011). This improved ability to identify errors may be derived from improved processing of sensory feedback during (or even before) the completion of the task (Beek & Lewbel, 1995). From this improved ability to identify errors, subsequent attempts can be corrected, thus improving success on actual performance of the task. Another theory suggests senior performers, or those subjects with greater familiarity or practice with the task, can monitor the motor commands sent to the body by comparison to a previously learned internal model of the expected sensory consequences of the movement (Wolpert & Ghahramani, 2000). Through this error detection capability, subjects can produce more effective motor behaviours in subsequent attempts. However, recognition of performance improvement may not always occur, as Alameddine and colleagues (Alameddine et al., 2018) showed that even though self- and objective evaluations of the suturing task improved with training, participants still underestimated their overall performance by half of the objective score objective evaluators assigned.

### Purpose of the task

Seven studies (Bested et al., 2019; Bolognini et al., 2016; Broman et al., 1992; Kennedy et al., 2014; Koppelaar & Wells, 2005; San Diego et al., 2014; Shirzad & Van der Loos, 2015) investigated a task that was novel to the participants, but trivial in nature (e.g., finger tapping, button pressing, or gripping tasks). Motor learning schema theory contends that the movement outcome of a novel motor task can be produced even if it had not been performed previously by the individual, because the parameters of the new task are based on the known performance of previously executed similar movements (Schmidt & Lee, 2011). Since gripping and button pressing are trivial tasks performed in everyday life (though perhaps not to the parameters defined by the experiments’ tasks), it stands to reason that a subject’s representation of the previously executed similar movements would help inform the self-evaluation of their performance on the novel motor task. That is, even though the parameters of the task are novel to the individual, they would not be guessing about the accuracy of their ability to perform the novel task, if they had a similar movement pattern to reference. Similar to how the cumulative amount of practice improves self-evaluation of performance, perhaps how closely related the novel task is to previously well-practiced movements influence the degree of accuracy of self-evaluation of that novel task.

The remaining sixteen studies (Alameddine et al., 2018; Alhusuny et al., 2020; Bao, Howard, et al., 2006; Casswell et al., 2016; Gao et al., 2019; Hadley et al., 2015; Hoozemans et al., 2001; Hu et al., 2015; Leopold et al., 2005; Mannella et al., 2019; Manzone et al., 2014; Nebiker et al., 2015; San Diego et al., 2014; Van Sickle et al., 2011; Ventimiglia et al., 2020; Vyasa et al., 2017; Wright et al., 2012) had participants perform a task directly related to their occupation or that for which they were training. The tasks were required for success in the profession and could also be considered high risk in that there was a negative outcome associated with poor performance of the task. For example, an adverse outcome of cataract surgery could result blindness. In such cases, learners may be more invested in identifying performance errors to ensure future success. Thus, these performers may have a greater tendency to perceive performance errors that may not exist objectively, resulting in an underestimation of performance. This may explain the results of 3 of the 6 surgical task studies (Hadley et al., 2015; Mannella et al., 2019; Nebiker et al., 2015) (which reported sufficient information to compare evaluations) whose participants underscored their performance. Additionally, how realistic the task is to one’s occupation may also mediate self-evaluation. For example, the self-evaluation scores of a task that more closely represented an actual cavity preparation (tooth in-jaw simulated model versus hand-held tooth only) had lower agreement with the objective-evaluation scores, and on average, participants underestimated their performance (San Diego et al., 2014). One possible interpretation is that when a medical task closely resembles the task to be performed on a patient, the participant becomes increasingly critical of their performance.

### Task difficulty

A study by Kennedy and coauthors, involving a computer task that required participants to move the mouse to intercept descending objects with the cursor (Kennedy et al., 2014). The task progressed from easy to very difficult trials that would undoubtedly cause object misses. Participants were asked to self-evaluate their performance (i.e., successful hit rate) after each trial. Interestingly, participants were less accurate and rated their performance poorer on the easier trials compared to the difficult ones. While their objective and self-evaluation of performance did decline for levels of increasing difficulty, participants’ accuracy of self-evaluation was greater at the moderate to most-difficult levels. Task difficulty expectations are related to interest in the task, with easy tasks eliciting greater interest than hard tasks (Hom & Maxwell, 1983).Therefore, one potential explanation is that participants may be more critical of their performance during an easy task for which they are paying greater attention. In other words, if one feels as though they made a mistake when they should not have, they score themselves much poorer than they otherwise would have. In contrast, when the difficulty level is high, errors in performance may be expected, resulting in a self-evaluation more in line with objective measures. Factors such as interest, attention, and motivation with respect to difficulty in motor performance self-evaluation may be avenues for future investigation.

### Task demonstration

Two studies included task demonstration as part of the study design and had contrasting results. Mannella and co-authors(Mannella et al., 2019) had surgery residents perform and self-evaluate five laparoscopic tasks, before and after observing a demonstration of correct performance of each task. They found that a demonstration did not influence self-evaluation. Conversely, Bested and colleagues (Bested et al., 2019) investigated the effect of a robotic demonstration on performance of a golf putting task in two groups of participants. One group received the demonstration prior to 50% of their trials and the other group received no demonstration. Participants evaluated their performance by estimating the ball endpoint location. Only the group that received demonstration significantly improved in both the accuracy and variability of the self-evaluation. In summary, demonstration prior to performance may improve self-evaluation of some types of motor tasks, but further investigation is necessary.

### Limitations

This systematic scoping review has some limitations. Studies published in languages other than English were not included due to the language limitations of the research team. Moreover, the motor tasks and task complexity, and evaluation measurement tools included in this review are heterogeneous which prevented a meta-analysis and limited our ability to draw definitive conclusions. Finally, this review was limited to upper-extremity tasks only and thus may not generalize to the self-evaluation of full-body or lower-extremity motor tasks such as balance and walking.

## Recommendations and conclusions

Future studies that investigate self-evaluation of motor performance as a primary objective may consider these recommendations. First, future studies should address the methodological issues identified in the included studies such as blinding. For example, if the study includes participants of different levels of experience or receive different types of feedback, objective expert assessors should be blinded to group allocation to reduce observer bias. Second, the definition of successful performance of the motor task should be clearly defined to the participant to ensure they attend to the aspect of their performance the investigators want evaluated. Asking a participant to self-evaluate with general questions such as ‘how did you perform?’, may lead to different interpretations of performance between individuals. Third, the self-evaluation response options should match the objective evaluation scoring criteria when possible, to improve the comparability and internal consistency. For instance, if an objective scoring domain is “time to completion” the self-evaluation tool should ask “how long did you take to perform this task?”. Finally, suggestions for future work include investigation of factors that may influence the accuracy of self-evaluation. These factors include degree of task difficulty, task novelty, perceived internal or external barriers to performance, the relatedness of the task to the performer’s training or profession and the consequences of task performance failure (or success). Additionally, future studies may want to investigate how these factors may apply to or differ in neurological conditions, such as stroke.

This review explored the relationship between self- and objective evaluation of upper-extremity task performance in healthy adults. We found that current research investigating this relationship as a primary study objective is limited. However, existing studies investigated this relationship as a secondary objective showed a moderate positive correlation between self- and objective evaluations, with individuals both under- and over-estimating performance. The accuracy of self-evaluation may depend on certain factors such as similarity between the evaluation methods, task familiarity and experience, practice, demonstration, and relatedness to real-life task situations. Rigorous studies are still needed to explain the accuracy of self-evaluation and the potential factors influencing self-evaluation of upper-extremity motor performance more clearly.

Acknowledgements, the authors would like to acknowledge Jessica Babineau, Information Specialist at Toronto Rehabilitation Institute, University Health Network, for her assistance in developing the search strategy and for conducting the database searches, and Dr. Dina Brooks, for her feedback and contributions to early drafts of this manuscript.

## Declaration of interest

none

## Funding source

This research was supported by a CIHR–NSERC collaborative grant (Fund # CPG-140198, CHRPJ478508-15).

---

Table 1. Type your title here.

Figure 1. Type your title here. Obtain permission and include the acknowledgement required by the copyright holder if a figure is being reproduced from another source.

## Notes

### Competing Interest Statement

The authors have declared no competing interest.

## References

Alameddine, M. B., Englesbe, M. J., & Waits, S. A. (2018). A Video-Based Coaching Intervention to Improve Surgical Skill in Fourth-Year Medical Students. Journal of Surgical Education, 1–5. https://doi.org/10.1016/j.jsurg.2018.04.003

Alhusuny, A., Cook, M., Khalil, A., Treleaven, J., Hill, A., & Johnston, V. (2020). Impact of accommodation, convergence and stereoacuity on perceived symptoms and surgical performance among surgeons. Surgical Endoscopy. https://doi.org/10.1007/s00464-020-08167-2

Bao, S., Howard, N., Spielholz, P., & Silverstein, B. (2006). Quantifying repetitive hand activity for epidemiological research on musculoskeletal disorders--part II: comparison of different methods of measuring force level and repetitiveness. Ergonomics, 49(4), 381–392. https://doi.org/10.1080/00140130600555938

Bao, S., Spielholz, P., Howard, N., & Silverstein, B. (2006). Quantifying repetitive hand activityfor epidemiological research on musculoskeletal disorders – part I: individual exposure assessment. Ergonomics, 49(4), 361–380. https://doi.org/10.1080/00140130500520214

Beek, P. J., & Lewbel, A. (1995). The science of juggling. Scientific American, 273, 92–97.

Bested, S. R., de Grosbois, J., Crainic, V. A., & Tremblay, L. (2019). The influence of robotic guidance on error detection and correction mechanisms. Human Movement Science, 66(December 2018), 124–132. https://doi.org/10.1016/j.humov.2019.03.009

Blakemore, S. J., Frith, C. D., & Wolpert, D. M. (2002). Abnormalities in the awareness and control of action. Trends in Cognitive Sciences, 6(6), 237–242. https://doi.org/10.1098/rstb.2000.0734

Bolognini, N., Zigiotto, L., Izzadora Souza Carneiro, M., & Vallar, G. (2016). “How Did I Make It?”: Uncertainty about Own Motor Performance after Inhibition of the Premotor Cortex Nadia. Journal of Cognitive Neuroscience, 28(7), 1052–1061. https://doi.org/10.1162/jocn

Broman, J.-E., Lundh, L.-G., Aleman, K., & Hetta, J. (1992). Subjective and Objective Performance in Patients with Persistent Insomnia. Scandinavian Journal of Behaviour Therapy, 21(3), 115–126. https://doi.org/10.1080/16506079209455903

Casswell, E. J., Salam, T., Sullivan, P. M., & Ezra, D. G. (2016). Ophthalmology trainees’ self-assessment of cataract surgery. British Journal of Ophthalmology, 100(6), 766–771. https://doi.org/10.1136/bjophthalmol-2015-307307

De Vet, H. C. W., De Bie, R. A., Van Der Heijden, G. J. M. G., Verhagen, A. P., Sijpkes, P., & Knipschild, P. G. (1997). Systematic reviews on the basis of methodological criteria. Physiotherapy, 83(6), 284–289. https://doi.org/10.1016/S0031-9406(05)66175-5

Estabrooks, C. A., Cummings, G. G., Olivo, S. A., Squires, J. E., Giblin, C., & Simpson, N. (2009). Effects of shift length on quality of patient care and health provider outcomes: systematic review. Quality and Safety in Health Care, 18(3), 181–188. https://doi.org/10.1136/qshc.2007.024232

Finez, L., Berjot, S., Rosnet, E., Cleveland, C., & Tice, D. M. (2012). Trait self-esteem and claimed self-handicapping motives in sports situations. Journal of Sports Sciences, 30(16), 1757–1765. https://doi.org/10.1080/02640414.2012.718089

Fourneret, P., & Jeannerod, M. (1998). Limited conscious monitoring of motor performance in normal subjects. Neuropsychologia, 36(11), 1133–1140. https://doi.org/10.1016/S0028-3932(98)00006-2

Gao, J., Liu, S., Feng, Q., Zhang, X., Jiang, M., Wang, L., Zhang, J., & Zhang, Q. (2019). Subjective and objective quantification of the effect of distraction on physician’s workload and performance during simulated laparoscopic surgery. Medical Science Monitor, 25, 3127–3132. https://doi.org/10.12659/MSM.914635

Guadagnoli, M. A., & Kohl, R. M. (2001). Knowledge of results motor learning: Relationship between error estimation and knowledge of results frequency. Journal of Motor Behaviour, 33(2), 217–224.

Hadley, C., Lam, S. K., Briceño, V., Luerssen, T. G., & Jea, A. (2015). Use of a formal assessment instrument for evaluation of resident operative skills in pediatric neurosurgery. Journal of Neurosurgery. Pediatrics, 16(November), 1–8. https://doi.org/10.3171/2015.1.PEDS14511

Henson, R. K. (2001). Understanding internal consistency reliability estimates: A conceptual primer on coefficient alpha. Measurement and Evaluation in Counseling and Development, 34(3), 177–189. https://doi.org/10.1080/07481756.2002.12069034

Hom, H. L., & Maxwell, F. R. (1983). The impact of task difficulty expectations on intrinsic motivation. Motivation and Emotion, 7(1), 19–24. https://doi.org/10.1007/BF00992962

Hoozemans, M. J. M., Beek, A. J. Van Der, Frings-dresen, M. H. W., & Molen, H. F. Van Der. (2001). Evaluation of methods to assess push / pull forces in a construction task. Applied Ergonomics, 32, 509–516.

Hu, Y., Kim, H., Mahmutovic, A., Choi, J., Le, I., & Rasmussen, S. (2015). Verification of accurate technical insight: a prerequisite for self-directed surgical training. Advances in Health Sciences Education : Theory and Practice, 20(1), 181–191. https://doi.org/10.1007/s10459-014-9519-3

Kennedy, P., Miele, D. B., & Metcalfe, J. (2014). The cognitive antecedents and motivational consequences of the feeling of being in the zone. Consciousness and Cognition, 30, 48–61. https://doi.org/10.1016/j.concog.2014.07.007

Kmet, L. M., Lee, R. C., & Cook, L. S. (2004). Standard Quality Assessment Criteria for Evaluating Primary Research Papers. Alberta Heritage Foundation for Medical Research, 13(February), 1–22.

Koppelaar, E., & Wells, R. (2005). Comparison of measurement methods for quantifying hand force. Ergonomics, 48(8), 983–1007. https://doi.org/10.1080/00140130500120841

Leopold, S. S., Morgan, H. D., Kadel, N. J., Gardner, G. C., Schaad, D. C., & Wolf, F. M. (2005). Impact of Educational Intervention on Confidence and Competence in the Performance of a Simple Surgical Task. The Journal of Bone & Joint Surgery, 87(5), 1031–1037. https://doi.org/10.2106/JBJS.D.02434

Liu, J., & Wrisberg, C. A. (1997). The effect of knowledge of results delay and the subjective estimation of movement form on the acquisition and retention. Research Quarterly for Exercise and Sport, 68(2), 145–151.

Mannella, P., Malacarne, E., Giannini, A., Russo, E., Caretto, M., Papini, F., Montt Guevara, M. M., Pancetti, F., & Simoncini, T. (2019). Simulation as tool for evaluating and improving technical skills in laparoscopic gynecological surgery. BMC Surgery, 19(1), 1–9. https://doi.org/10.1186/s12893-019-0610-9

Manzone, J., Tremblay, L., You-Ten, K. E., Desai, D., & Brydges, R. (2014). Task-versus ego-oriented feedback delivered as numbers or comments during intubation training. Medical Education, 48(4), 430–440. https://doi.org/10.1111/medu.12397

McHugh, M. L. (2012). Interrater reliability: the kapa statistic. Biochemia Medica, 22(3), 276–282.

Moher, D., Liberati, A., Tetzlaff, J., Altman, D. G., & Group, P. (2009). Preferred reporting items for systematic reviews and meta-analyses: the PRISMA statement. PLoS Medicine, 6(7), e1000097.

Mukaka, M. M. (2012). A guide to appropriate use of Correlation coefficient in medical research. Malawi Medical Journal, 24(3), 69–71. https://doi.org/10.1016/j.cmpb.2016.01.020

Nebiker, C. A., Mechera, R., Rosenthal, R., Thommen, S., Marti, W. R., von Holzen, U., Oertli, D., & Vogelbach, P. (2015). Residents’ performance in open versus laparoscopic bench-model cholecystectomy in a hands-on surgical course. International Journal of Surgery, 19, 15–21. https://doi.org/10.1016/j.ijsu.2015.04.072

Peters, M. D. J., Godfrey, C. M., Khalil, H., McInerney, P., Parker, D., & Soares, C. B. (2015). Guidance for conducting systematic scoping reviews. International Journal of Evidence-Based Healthcare, 13(3), 141–146. https://doi.org/10.1097/XEB.0000000000000050

San Diego, J. P., Newton, T., Quinn, B. F. A., Cox, M. J., & Woolford, M. J. (2014). Levels of agreement between student and staff assessments of clinical skills in performing cavity preparation in artificial teeth. European Journal of Dental Education, 18(1), 58–64. https://doi.org/10.1111/eje.12059

Schmidt, R. A. (2003). Motor schema theory after 27 years: Reflections and implications for a new theory. Research Quarterly for Exercise and Sport, 74(4), 366–375. https://doi.org/10.1080/02701367.2003.10609106

Schmidt, R. A., & Lee, T. D. (2011). Motor control and learning: a behavioral emphasis (5th ed.). Human Kinetics.

Shepperd, J. A., & Arkin, R. M. (1990). Shyness and self-presentation. In W. R. Crozier (Ed.), Shyness and embarrassment - Perspectives from social psychology. Cambridge University Press.

Shirzad, N., & Van Der Loos, H. F. M. (2012). Error amplification to promote motor learning and motivation in therapy robotics. Proceedings of the Annual International Conference of the IEEE Engineering in Medicine and Biology Society, EMBS, March, 3907–3910. https://doi.org/10.1109/EMBC.2012.6346821

Shirzad, N., & Van der Loos, H. F. M. (2015). Evaluating the User Experience of Exercising Reaching Motions With a Robot That Predicts Desired Movement Difficulty. Journal of Motor Behaviour, 48(1), 31–46. https://doi.org/10.1080/00222895.2015.1035430

Squires, J. E., Estabrooks, C. A., Gustavsson, P., & Wallin, L. (2011). Individual determinants of research utilization by nurses: a systematic review update. Implement Sci, 6, 1. https://doi.org/10.1186/1748-5908-6-1

Van Sickle, K. R., Buck, L., Willis, R., Mangram, A., Truitt, M. S., Shabahang, M., Thomas, S., Trombetta, L., Dunkin, B., & Scott, D. (2011). A multicenter, simulation-based skills training collaborative using shared GI mentor II systems: Results from the Texas association of surgical skills laboratories (TASSL) flexible endoscopy curriculum. Surgical Endoscopy and Other Interventional Techniques, 25(9), 2980–2986. https://doi.org/10.1007/s00464-011-1656-7

Ventimiglia, E., Sindhubodee, S., Besombes, T., Pauchard, F., Quadrini, F., Delbarre, B., Jiménez Godínez, A., Barghouthy, Y., Corrales Acosta, M. A., Kamkoum, H., Villa, L., Doizi, S., Somani, B. K., & Traxer, O. (2020). Operator-assisted vs self-achieved basketing during ureteroscopy: results from an in vitro preference study. World Journal of Urology. https://doi.org/10.1007/s00345-020-03431-5

Vyasa, P., Willis, R. E., Dunkin, B. J., & Gardner, A. K. (2017). Are General Surgery Residents Accurate Assessors of Their Own Flexible Endoscopy Skills? Journal of Surgical Education, 74(1), 23–29. https://doi.org/10.1016/j.jsurg.2016.06.018

Wolpert, D. M., & Ghahramani, Z. (2000). Computational principles of movement neuroscience. Nature Neuroscience, 3(11s), 1212–1217. https://doi.org/10.1038/81497

Wright, A. S., McKenzie, J., Tsigonis, A., Jensen, A. R., Figueredo, E. J., Kim, S., & Horvath, K. (2012). A structured self-directed basic skills curriculum results in improved technical performance in the absence of expert faculty teaching. Surgery (United States), 151(6), 808–814. https://doi.org/10.1016/j.surg.2012.03.018

